# A gap-free tomato genome built from complementary PacBio and Nanopore long DNA sequences reveals extensive linkage drag during breeding

**DOI:** 10.1101/2021.08.30.456472

**Authors:** Willem M. J. van Rengs, Maximilian H.-W. Schmidt, Sieglinde Effgen, Yazhong Wang, Mohd Waznul Adly Mohd Zaidan, Bruno Huettel, Henk J. Schouten, Björn Usadel, Charles J. Underwood

**Affiliations:** Department of Chromosome Biology, Max Planck Institute for Plant Breeding Research, Carl-von-Linné-Weg 10, 50829, Cologne, Germany; IBG-4 Bioinformatics, Forschungszentrum Jülich, 52428 Jülich, Germany; Max Planck-Genome-center Cologne, Carl-von-Linné-Weg 10, 50829 Cologne, Germany; Department of Plant Breeding, Wageningen University and Research, P.O. Box 386, 6700 AJ, Wageningen, The Netherlands; Heinrich Heine University Düsseldorf, Institute of Biological Data Science, Düsseldorf, Germany

**Keywords:** Tomato, PacBio, SMRT sequencing, Oxford Nanopore Technologies, Nanopore sequencing, Genome assembly, Linkage drag, Plant breeding, Tobacco mosaic virus, Moneyberg-TMV

## Abstract

The assembly and scaffolding of plant crop genomes facilitates the characterization of genetically diverse cultivated and wild germplasm. The cultivated tomato has been improved through the introgression of genetic material from related wild species, including resistance to pandemic strains of Tobacco Mosaic virus (TMV) from *Solanum peruvianum*. Here we applied PacBio HiFi and ONT nanopore sequencing to develop independent, highly contiguous and complementary assemblies of an inbred TMV-resistant tomato variety. We merged the HiFi and ONT assemblies to generate a long-read-only assembly where all twelve chromosomes were represented as twelve contiguous sequences (N50=68.5 Mbp). The merged assembly was validated by chromosome conformation capture data and is highly consistent with previous tomato assemblies that made use of genetic maps and HiC for scaffolding. Our long-read-only assembly reveals that a complex series of structural variants linked to the TMV resistance gene likely contributed to linkage drag of a 64.1 Mbp region of the *S. peruvianum* genome during tomato breeding. We show that this minimal introgression region is present in six cultivated tomato hybrid varieties developed in three commercial breeding programs. Our results suggest that complementary long read technologies can facilitate the rapid generation of near complete genome sequences.

## Introduction

DNA sequencing technology has evolved rapidly in the last two decades, and long read DNA sequencing has fundamentally altered approaches in genome assembly. Long DNA sequence reads that can span repeated sequences, including the full length of transposable elements, can simplify the assembly process (Koren *et al*., 2012; Dumschott *et al*., 2020). Pacific Biosciences (PacBio) initially delivered long yet error prone reads (initially with a raw base accuracy of 82.1 – 84.6%) facilitating short-long read hybrid assembly and long-read-only assembly approaches (Koren *et al*., 2012; Berlin *et al*., 2015). Whilst this was also rapidly adopted in the field of plant genomics (Vanburen *et al*., 2015; Zapata *et al*., 2016), the application of nanopore sequencing developed by Oxford Nanopore Technologies (ONT) led to long read sequencing with minimal capital outlay (Schmidt *et al*., 2017; Michael *et al*., 2018). Increases in sequencing yield and improvements in raw base accuracy, as well as protocol development for DNA extraction (Vaillancourt and Buell, 2019; Vilanova *et al*., 2020), have facilitated the analysis of crop pangenomes (Alonge *et al*., 2020; Liu *et al*., 2020; Qin *et al*., 2021), and the development of near-complete human and Arabidopsis genome sequences (Nurk *et al*., 2021; Naish *et al*., 2021).

The PacBio Sequel II platform now delivers long High Fidelity (HiFi) reads with a low error rate. HiFi reads are a consensus of multiple passes of the same circular DNA molecule, typically with a 10-20 kbp genomic DNA insert, and therefore have a much higher per base accuracy (99.5 - 99.9%) than the individual passes (85%) (Vollger *et al*., 2020; Hon *et al*., 2020). The length and low error rate of HiFi reads means that they span the majority of repeated sequences and can also differentiate between highly similar repeated sequences. Various assembly tools have been developed to harness the characteristics of HiFi reads (Nurk *et al*., 2020; Cheng *et al*., 2021). HiFi reads are restricted in length because larger DNA insert sizes are sequenced with a lower number of passes, due to limitations of DNA polymerase processivity. ONT does not currently support multiple pass consensus sequencing and has largely focused on applying deep learning to improve raw read base accuracy to 97% and the forthcoming Q20+ chemistry promises 99% accuracy and potentially higher. ONT sequencing does not yet provide a raw base accuracy high enough to use the same assembly tools that are used for HiFi reads, yet can generate reads that are multiple megabases in length that can reliably align to reference genomes (Payne *et al*., 2019). ONT reads in the 20 kbp - 200 kbp category can span complex tandem arrays of repeated sequences that litter plant genomes. Therefore, current long read sequencing options are either very high-quality reads between the length of 10-20 kbp (HiFi) or considerably longer reads with lower per base accuracy (ONT). Despite the apparently complementary strengths of HiFi and ONT data, there have been few attempts to combine the two data types in genome assemblies.

Tomato (*Solanum lycopersicum*) is the world’s most popular fruit and an important source of vitamins and minerals in a balanced diet. Tomato varieties can be improved by the introgression of novel resistances to abiotic and biotic stresses from multiple related wild species (Foolad *et al*., 1997; Bai and Lindhout, 2007; Peralta *et al*., 2008; Li *et al*., 2010). High quality reference genomes are available for the cultivated tomato, *S. lycopersicum* cv. ‘Heinz 1706’ (SL4.0) (Sato *et al*., 2012; Hosmani *et al*., 2019), the closely related red-fruited *S. pimpinellifolium* ‘LA2093’ (Wang *et al*., 2020*b*), the more distantly related green fruited *S. pennellii* ‘LA0716’ (Bolger *et al*., 2014) and the purple/black fruited *S. lycopersicoides* ‘LA2951’ (Powell *et al*., 2020). Alongside these reference genomes, more than 1000 tomato varieties and wild tomato species have been sequenced on Illumina platforms (Aflitos *et al*., 2014; Lin *et al*., 2014; Gao *et al*., 2019) and a panel of 100 of these were recently sequenced using ONT long reads to characterize structural variants (Alonge *et al*., 2020). In summary, cultivated and wild tomato genomes have been extensively characterized to understand genetic diversity yet even the SL4.0 reference genome contains gaps in repetitive regions, including subtelomeric and centromeric regions.

Tobamoviruses are single stranded RNA viruses that can be devastating pathogens for tomato and other vegetable crops. The tobacco mosaic virus (TMV) and tomato mosaic virus (ToMV) are two tobamoviruses that can infect susceptible tomato cultivars and lead to substantial yield reduction (Lanfermeijer *et al*., 2003, 2005). From the 1930s until the 1960s tomato breeders searched for TMV resistance in various wild tomato species, including *S. pimpinellifolium, S. pennellii, S. habrochaites* (syn. *L. hirsutum*), *S. chilense* and *S. peruvianum* (Pelham, 1966). TMV resistance was found including in *S. pennellii, S. habrochaites* (*Tm-1*) and *S. peruvianum* (allelic resistance genes *Tm-2* and *Tm-2*^*2*^) (Pelham, 1966; Lanfermeijer *et al*., 2005). Of the three known TMV resistance genes, introgression of *Tm-2*^*2*^ from *S. peruvianum* (accession P. I. 128650) has provided dominant, robust and lasting resistance to all TMV and ToMV strains since its discovery in the 1960s and therefore most modern tomato greenhouse varieties contain the *Tm-2*^*2*^ gene (Alexander, 1963; Schouten *et al*., 2019). The *Tm-2*^*2*^ gene encodes a coiled-coil domain Nod-like receptor (CC-NLR) protein that recognizes the movement protein (MP), a key viral protein that facilitates TMV/ToMV movement between plant cells via plasmodesmata (Lanfermeijer *et al*., 2003, 2005; Wang *et al*., 2020*a*). The exact size and the structure of the introgression containing the *Tm-2*^*2*^ gene has remained unknown, although it was known to include at least half of the physical length of chromosome 9 (Lin *et al*., 2014; Schouten *et al*., 2019).

Here we sequenced and assembled the genome of the *S. lycopersicum* cultivar Moneyberg-TMV (see Materials and methods for breeding history), a popular line used in tomato research because of its vigour, homozygosity and capacity for genetic transformation. PacBio HiFi and ONT nanopore sequencing reads were independently used to develop highly contiguous genome assemblies. We merged HiFi and ONT assemblies to develop a long-read-only assembly (‘MbTMV’) where all twelve chromosomes were present as single contigs without the introduction of artificial gaps. The MbTMV assembly was validated through orthogonal approaches including chromosome conformation capture data, mapping of raw long read data and coverage analysis, and extensive comparison with the SL4.0 assembly. Through our long-read-only assembly and the analysis of six commercial tomato hybrids, we show that a minimal 64.1 Mbp introgression has been dragged along with the *Tm-2*^*2*^ gene despite more than fifty years of tomato breeding.

## Results

We generated long read DNA sequences of the *S. lycopersicum* cv. ‘Moneyberg-TMV’ genome using two different third generation sequencing technologies. High molecular weight (HMW) DNA was extracted and used to construct HiFi sequencing libraries where we made use of two different insert sizes to balance read length and read quality. The two libraries were subjected to PacBio HiFi sequencing on two Sequel II SMRT cells. The first cell yielded 18.8 Gbp with a HiFi read length N50 of 13.9 kbp and mean quality (Q-value) of 28.59 (Table 1). The second cell yielded 21.8 Gbp of data with a HiFi read length N50 of 24.8 kbp and mean Q-value of 24.65 (Table 1). Using the same HMW DNA two identical nanopore library preparations were performed and subjected to ONT long read sequencing on two PromethION cells. This yielded 100.5 and 101.1 Gbp with respective read length N50 of 41.6 and 42.4 kbp, and mean Q-values of 11.28 and 11.18 (Table 1).

**Table 1.** Statistics of different sequencing methods. The table shows the total sequencing output and generic summary statistics for two cells of PacBio HiFi data and two cells of ONT PromethION data, before quality filtering.

|  | PacBio |  |  | Oxford Nanopore Technologies |  |  |
| --- | --- | --- | --- | --- | --- | --- |
|  | Run_1 | Run_2 | Combined | Run_1 | Run_2 | Combined |
| Number of sequences | 1,356,407 | 867,228 | 2,223,635 | 4,597,421 | 4,522,199 | 9,119,620 |
| Output (Gbp) | 18.8 | 21.8 | 40.6 | 100.5 | 101.1 | 201.6 |
| Longest read (kbp) | 31.3 | 50.3 | 50.3 | 862.3 | 629.5 | 862.3 |
| Median read length (kbp) | 13.9 | 24.3 | 14.7 | 13.1 | 13.6 | 13.4 |
| Mean read length (kbp) | 13.9 | 25.1 | 18.3 | 21.9 | 22.4 | 22.1 |
| N50 read length (kbp) | 13.9 | 24.8 | 21.1 | 41.6 | 42.4 | 42.0 |
| Mean quality score | 28.59 | 24.65 | 26.06 | 11.28 | 11.18 | 11.23 |

We used four genome assembly tools to assemble the HiFi and ONT data, and compared the respective outcomes. PacBio HiFi reads were independently assembled using Hifiasm and Canu, with respective N50 values of 31.3 and 17 Mbp (Table 2). The L50 values were 10 (Hifiasm) and 14 (Canu), and both assemblies contained single contigs that spanned the full length of the SL4.0 reference chromosome 5 (Supplementary Fig. 1). We checked for the completeness of the assembly of gene sequences by Benchmarking Universal Single Copy Orthologs (BUSCO) analysis which showed both assemblies were almost gene complete (both 98.4%), slightly improving upon the SL4.0 genome and comparable to the *S. pimpinellifolium* ‘LA2093’ genome (Table 2 & 3). ONT reads were independently assembled using Flye and NECAT, with respective N50 values of 18.9 and 50.2 Mbp. The L50 values were 15 (Flye) and 7 (NECAT), and the NECAT assembly contained single contigs that spanned the length of SL4.0 chromosomes 5 and 7 (Table 2 & Supplementary Fig. 1). While the NECAT ONT assembly was more contiguous than either of the HiFi assemblies, gene completeness was lower in both the raw Flye (97.8%) and NECAT (97.1%) assemblies (Table 2), but polishing was able to improve these values. All four assemblies were aligned against the SL4.0 genome and were found to be collinear, irrespective of technology or assembly tool used, except for chromosome 9 where the *S. peruvianum* introgression resides, indicative of mostly correct assemblies (Supplementary Fig. 1).

**Table 2.** Statistics of different assembly methods. Table 2 shows the summary statistics for assemblies obtained from PacBio and ONT data, respectively. The Flye + NECAT merged assembly presented here does not include polishing of the merged assembly with PacBio data (see Materials and methods). BUSCO analysis of the completeness of gene content is using the *Solanales* benchmark set.

| Input data | 40.6 Gbp (PacBio HiFi) |  |  | 165.7 Gbp (ONT) |  |  |
| --- | --- | --- | --- | --- | --- | --- |
| Assembly | Hifiasm | Canu | Hifiasm + Canu | Flye | NECAT | Flye + NECAT |
| Number of contigs | 750 | 2396 | 731 | 371 | 125 | 105 |
| Cumulative size (Mbp) | 868.9 | 928.7 | 871.9 | 809.7 | 829.6 | 829.1 |
| N50 (Mbp) | 31.3 | 17.0 | 36.8 | 18.9 | 50.2 | 53.4 |
| N90 (Mbp) | 8.7 | 0.08 | 9.1 | 5.4 | 10.6 | 18.6 |
| L50 | 10 | 14 | 9 | 15 | 7 | 7 |
| L90 | 32 | 170 | 31 | 45 | 21 | 15 |
| Longest contig (Mbp) | 66.6 | 66.7 | 66.8 | 47.0 | 74.2 | 82.8 |
| BUSCO (genes searched) | 5950 | 5950 | 5950 | 5950 | 5950 | 5950 |
| BUSCO (complete) | 5852 | 5853 | 5852 | 5820 | 5779 | 5824 |
| BUSCO (complete single) | 5739 | 5735 | 5739 | 5703 | 5666 | 5713 |
| BUSCO (complete duplicated) | 113 | 118 | 113 | 117 | 113 | 111 |
| BUSCO (fragmented) | 12 | 12 | 12 | 21 | 38 | 17 |
| BUSCO (missing) | 86 | 85 | 86 | 109 | 133 | 109 |
| LAI | 13.48 | 9.65 | 13.43 | 10.63 | 10.39 | 10.58 |
| raw LAI | 7.85 | 4.02 | 7.80 | 5.00 | 4.76 | 4.95 |

**Table 3.** Statistics of different scaffolded assemblies. Table 3 shows the summary statistics for the MbTMV assembly, as well as for three “gold standard” tomato species genome assemblies: ‘Heinz 1706’ SL4.0 (Hosmani *et al*., 2019), *S. pimpinellifolium* ‘LA2093’ (Wang *et al*., 2020*b*) *and S. pennellii* ‘LA0716’ (Bolger *et al*., 2014). BUSCO analysis of the completeness of gene content is using the *Solanales* benchmark set.

| Assembly name | MbTMV | SL4.0 | LA2093 | LA0716 |
| --- | --- | --- | --- | --- |
| PacBio assembly | HiFiasm v0.14.2;<br>Canu v2.1.1 | Canu v1.5 | Canu v1.7.1 | - |
| ONT assembly | NECAT ; Flye | - | - | - |
| Polishing | Medaka (ONT);<br>Racon (HiFi) | PacBio +<br>Illumina | PacBio +<br>Illumina | Illumina |
| Scaffolding | QuickMerge | HiC | HiC | Genetic map |
| Number of scaffolds | 13 | 13 | 13 | 13 |
| Scaffolds<br>>50 Mbp | 12 | 11 | 11 | 13 |
| Cumulative size<br>(Mbp) | 833.0 | 782.5 | 810.5 | 989.5 |
| N50 (Mbp) | 68.5 | 65.3 | 67.2 | 78 |
| N90 (Mbp) | 54.7 | 53.5 | 55.6 | 60.7 |
| L50 | 6 | 6 | 6 | 6 |
| L90 | 11 | 11 | 11 | 12 |
| Maximum contig<br>size (Mbp) | 96.5 | 90.9 | 95.5 | 109.3 |
| Ns per 100 kbp | 0.64 | 5.72 | 357.21 | 11533.09 |
| BUSCO (genes<br>searched) | 5950 | 5950 | 5950 | 5950 |
| BUSCO (complete) | 5853 | 5825 | 5854 | 5847 |
| BUSCO (complete<br>single) | 5743 | 5718 | 5744 | 5711 |
| BUSCO (complete<br>duplicated) | 110 | 107 | 110 | 136 |
| BUSCO<br>(fragmented) | 12 | 24 | 11 | 17 |
| BUSCO (missing) | 85 | 101 | 85 | 86 |
| LAI | 13.43 | 10.39 | 13.18 | 7.70 |
| raw LAI | 7.80 | 4.76 | 7.55 | 2.07 |

To test whether sequencing coverage was saturating, we randomly downsampled the HiFi and ONT data sets in intervals of 10% (from 90% down to 10%). Subsequently, we performed 9 Hifiasm and 9 NECAT assemblies on the respective downsampled datasets (Supplementary Table 3 & 4). For Hifiasm, we found that downsampling to as little as 50% (20.3 Gbp; about 24x coverage) of the full HiFi dataset only had a marginal effect on contiguity, indicating that the coverage was saturating (Supplementary Table 3). For NECAT, which internally selects the longest 40x coverage for assembly, we noticed that downsampling to even just 30% (49.7 Gbp; about 60x coverage) again only had a marginal effect on contiguity. However, care must be taken to avoid NECAT assembly artifacts that occasionally happen even at high coverage (Supplementary Table 4).

The dotplots of the HiFi and ONT assemblies revealed partial complementarity of the assemblies as the breakpoints were different (Supplementary Fig. 1 & Supplementary Fig. 2), suggesting genome merging could further improve contiguity. We independently merged both ONT assemblies and both HiFi assemblies, followed by a subsequent merging of the two merged assemblies. In the final merged assembly, henceforth referred to as “MbTMV”, 98.9% of the assembly was represented in 12 contigs (N50=68.5 Mbp), all with a length of at least 52.2 Mbp, strongly suggestive of a near complete contig assembly (Table 3). Strikingly, when the MbTMV assembly was aligned against the SL4.0 genome assembly the twelve tomato chromosomes were represented as twelve contigs (Fig. 1a), as expected from chromosome counting (Fig. 1b). 8.56 Mbp of unplaced sequence was concatenated together on chromosome 0 and contained just 0.08% (5/5853) of BUSCO gene content identified in the MbTMV assembly, further demonstrating the twelve contig assembly is near complete.

**Fig. 1:**
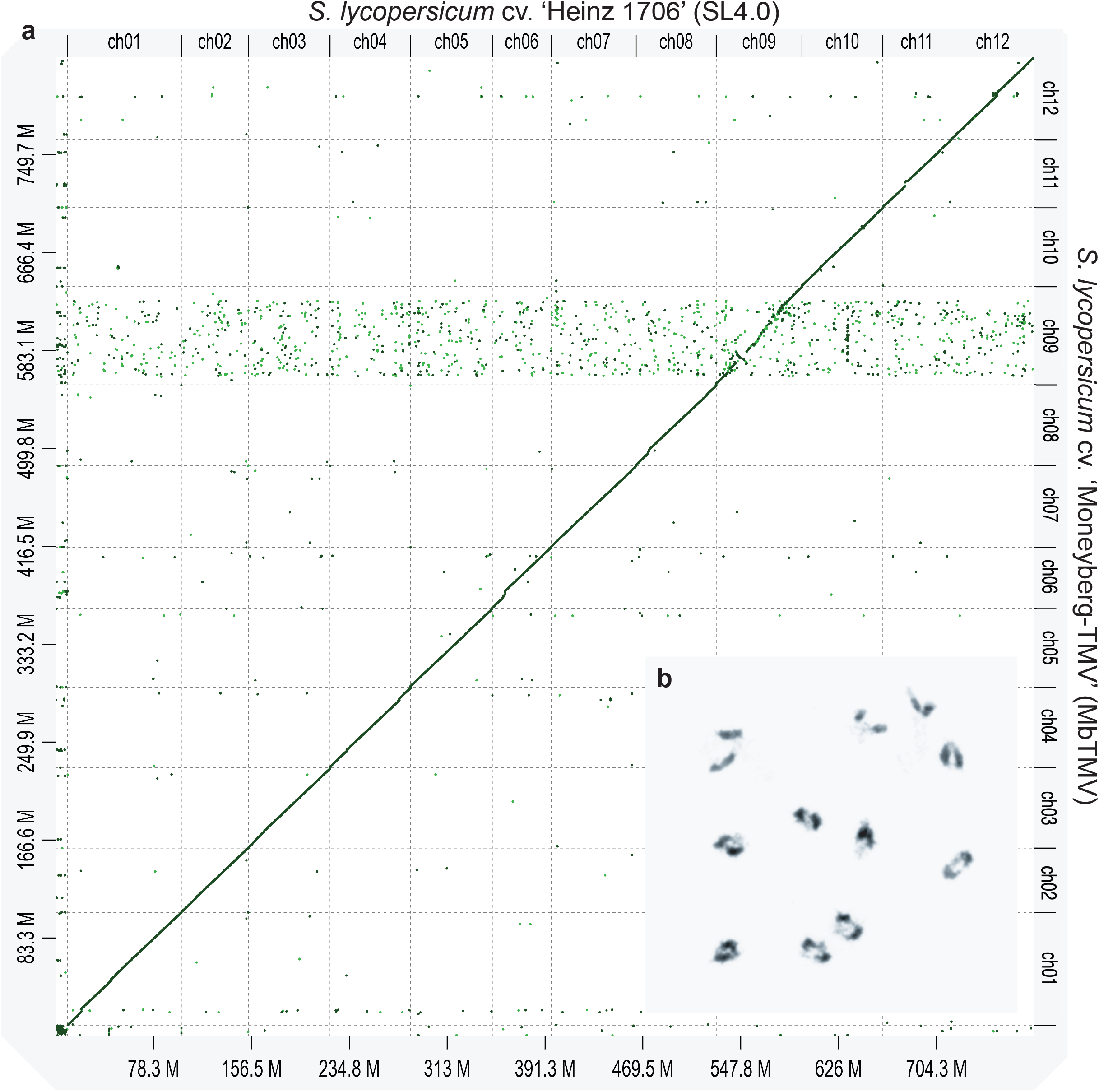
Alignment of the Moneyberg-TMV (MbTMV) assembly with Heinz 1706 (SL4.0) assembly. **a**, D-GENIES alignment of the MbTMV assembly against the SL4.0 assembly. Chromosome names were shortened to “ch00-ch12” for convenience, min identity (abs) was set to 0.65. All chromosomes show near perfect alignment except for ch09. The x-axis represents the potition on the SL4.0 genome, where M is megabase pairs. **b**, The 12 chromosomes of Moneyberg-TMV paired during meiosis, more specifically during diakinesis I.

In order to check the quality of the MbTMV assembly we made use of orthogonal sequencing and analysis approaches. Firstly, we generated high resolution Omni-C (chromosome conformation capture) data and aligned the paired reads to the MbTMV assembly, and assessed the resulting contact map. The contact map was supportive of well assembled chromosomes, with eleven of the twelve major contigs remaining unchanged (Fig. 2a). The single recommended change was the removal of 0.59 Mbp of repeat dense sequence at the end of chromosome 11, however the non-repetitive part of this region does align well to SL4.0 so we decided to maintain this region in our assembly (Supplementary Fig. 1 & Supplementary Fig. 3). Secondly, reference-based scaffolding of the individual HiFi and ONT assemblies (using the SL4.0 reference) led to assemblies consistent with the merged MbTMV assembly, whereas reference-based scaffolding of the merged MbTMV assembly itself did not improve or break the assembly (Table 3 & Supplementary Table 1). Thirdly, we remapped HiFi and ONT reads back to the MbTMV assembly to assess read coverage across the assembly (Fig. 2b). Indicative of an almost complete assembly, we found mostly even coverage across all twelve chromosomes, except in a few regions that are notoriously difficult to assemble. Higher ONT coverage was found on chromosome 1 and 2 overlapping with the highly repetitive 5S rDNA and 45S rDNA regions, respectively (Fig. 2b and Supplementary Fig. 5) (Schmidt-Puchta *et al*., 1989; Perry and Palukaitis, 1990; Zhong *et al*., 1998; Chang *et al*., 2008). A peak of HiFi and ONT coverage on chromosome 11 corresponded to a partial insertion of the mitochondrial genome, that has also been reported in the SL2.50 and SL4.0 assemblies (Fig. 2b) (Kim and Lee, 2018; Hosmani *et al*., 2019). Lower than average HiFi and ONT coverage was found adjacent to centromeric TRG4 dense regions on chromosomes 6, 8 and 11, likely due to difficulties in assembling tandem arrays of 45S rDNA derived satellite repeats that are known to occur in three clusters in the tomato genome (Fig. 2b) (Jo *et al*., 2009). However, other complete chromosomes (including 3, 5, 7, 9 and 10) had very stable coverage even over centromeric regions, indicating near complete assemblies. Fourthly, we were able to find clusters of the subtelomeric satellite sequence TGR1 in all subtelomeric regions apart from one end of chromosome 1 and both ends of chromosome 2, reflecting published FISH results (Fig. 2b)(Zhong *et al*., 1998). We also found long tandem arrays of the telomeric repeat ((TTTAGGG)_n_), on 14 of the 24 chromosome ends (Fig. 2b) (Ganal *et al*., 1991). Hifiasm (HiFi), Canu (HiFi) and NECAT all led to near identical assemblies of tandemly repeated sequences in the centromere of chromosome 5, which was also represented in the final MbTMV merged assembly, adding confidence that the centromere had been spanned in full (Fig. 2e, Fig. 2f & Supplementary Fig. 4). Fifthly, chromosome lengths were compared with three “gold standard” tomato species genomes (Fig. 2c). MbTMV had slightly longer chromosomes than the SL4.0 and *S. pimpinellifolium* ‘LA2093’ genomes in all cases except for chromosome 2. Additionally, the MbTMV genome had the least unplaced sequences and the lowest number of Ns per 100 kbp of all four genomes analysed (all Ns in MbTMV are found on the small chromosome 0)(Fig. 2c & Table 3). We suspected this could be related to a better assembly of repeat content, including the subtelomeres and telomeres, but also Long Terminal Repeat (LTR) transposable elements found in pericentromeric regions. We therefore independently assessed the genomes for completeness of LTRs using the LTR Assembly Index (LAI) metric (Ou *et al*., 2018). We found the MbTMV assembly had the highest LAI value, suggesting repeats were better assembled, which may partially explain longer chromosome lengths (Fig. 2c & Table 3). The MbTMV chromosome 9 was roughly 15.25 Mb longer than that from SL4.0 and similar in size to that from *S. pennellii* ‘LA0716’ (Fig. 2c). Finally, we compared MbTMV with SL4.0 using Synteny and Rearrangement Identifier (SyRI) (Goel *et al*., 2019) and identified syntenic regions, SNPs, insertions, deletions and other structural variations such as duplications, translocations and inversions (Fig. 2d & Supplementary Table 5). Highly increased variation is observed on chromosome 9, where the *S. peruvianum* introgression is located, with a series of large inversions and extended non-syntenic regions (Fig. 2d & Supplementary Table 5). All other chromosomes were highly syntenic between the MbTMV assembly and SL4.0, except for a number of duplications that were identified (Fig. 2d). In summary, the MbTMV genome sequence represents a near complete assembly of all tomato genes and nuclear chromosomes.

**Fig. 2:**
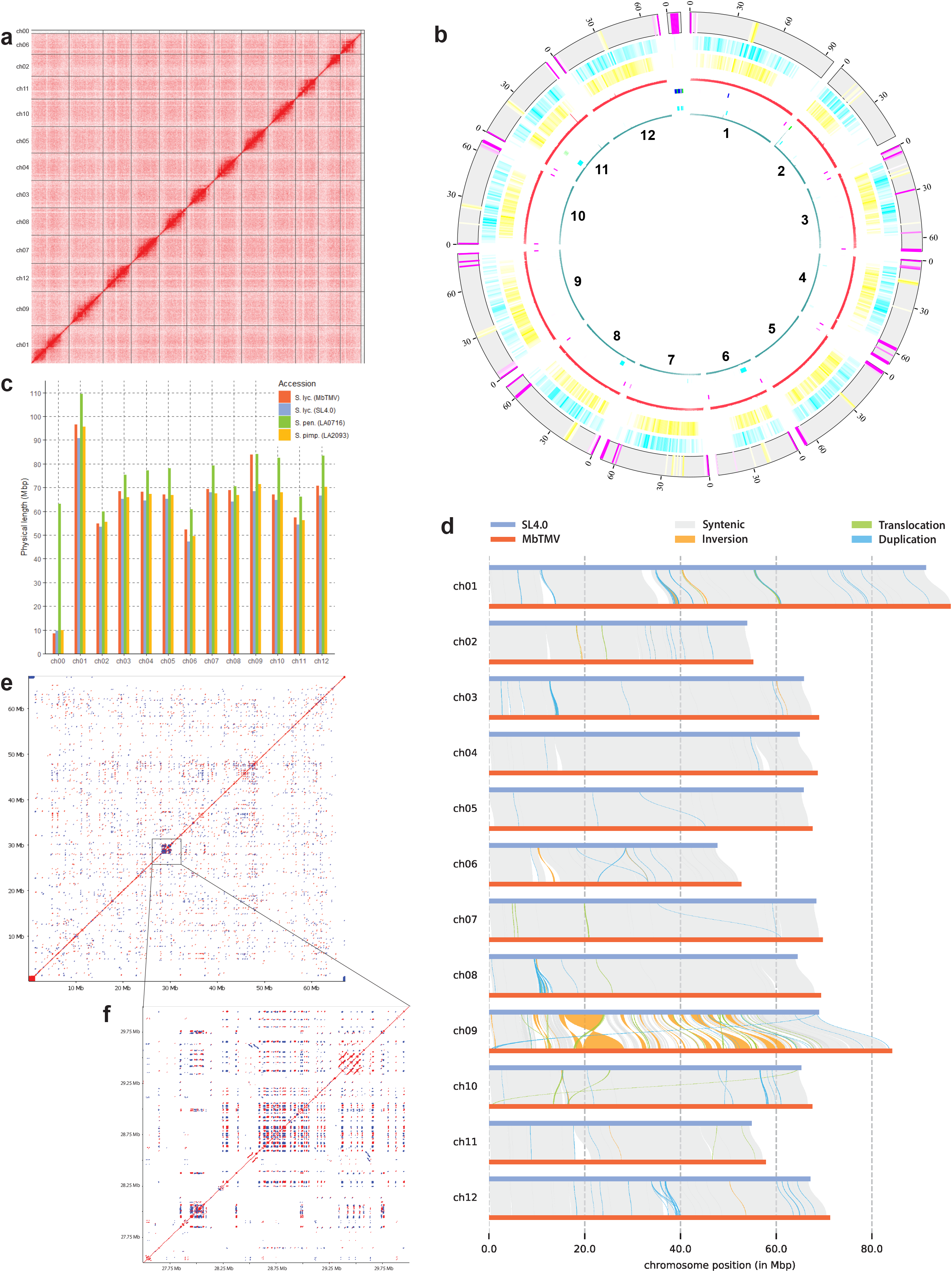
Validation of MbTMV assembly. **a**, Omni-C validation (Hi-C plot as plotted by Juicebox) demonstrating strong interactions within each chromosome of MbTMV. **b**, Circos plot of the MbTMV assembly including coverage of raw reads and genomic elements. From inner to outer ring: Ring 1 - ONT read coverage (cyan); Ring 2 - 45S rDNA intergenic spacer (IGS) (light blue); Ring 3 - telomeric repeat (pink), 5S rDNA sequence (dark blue), 45S rDNA (light green); Ring 4 – PacBio HiFi read coverage (red); Ring 5 – TGR3 (yellow); Ring 6 – TGR2 (light blue); Ring 7 – TGR1 (pink), TGR4 centromeric repeat (yellow) **c**, Chromosome lengths plot of MbTMV assembly including previously published assemblies: *S. lycopersicum* cv. ‘Heinz 1706’ (SL4.0), *S. pennellii* ‘LA0716’ and *S. pimpinellifolium* ‘LA2093’. **d**, Synteny and Rearrangement (SyRI) plot of all 12 MbTMV chromosomes (Query) against all 12 SL4.0 chromosomes (Ref). All chromosomes show high synteny except for chromosome 9. Non-syntenic (non-matching) regions are shown as white gaps between Ref and Query. **e**, Self alignment and dot plot analysis of the full length of MbTMV chromosome 5. **f**, Self alignment and dot plot analysis of the centromeric region of MbTMV chromosome 5.

The *Tm-2*^*2*^ gene confers broad resistance to TMV and ToMV and is located within a large introgression from *S. peruvianum* on chromosome 9, yet the full sequence and structure of the region is not known (Lin *et al*., 2014). We re-aligned chromosome 9 sequences from MbTMV and SL4.0, called variants and used polymorphism density to map the exact break points of the introgression. This revealed that the introgression from *S. peruvianum* is 64.1 Mbp (Fig 3a, Supplementary Fig. 6 & Supplementary Table 6). The *S. peruvianum* sequence contains several large inversions compared to SL4.0, but the *Tm-2*^*2*^ gene is located within a syntenic region (Fig. 3a). To better understand the local context of the *Tm-2*^*2*^ gene in the fully assembled chromosome, we performed further alignments between MbTMV and SL4.0 sequences from ∼10 Mbp and ∼1 Mbp regions centered upon the *Tm-2*^*2*^ gene (Fig. 3b & Fig. 3c). Immediately at the start of the introgression, a drop in synteny is observed due to a series of insertions in the *S. peruvianum* derived sequence, followed by several large insertions and deletions, and two large inversions (Fig. 3b & Supplementary Table 7). At the local scale, the *Tm-2*^*2*^ gene is located within a syntenic region of about 150.5 kbp that is immediately adjacent to a non-syntenic region that is estimated to be 101.7 kbp longer in *S. peruvianum* (Fig. 3c & Supplementary Table 8). We validated our chromosome 9 assembly by aligning ONT reads from a different inbred variety (LYC 1969) that contains the *Tm-2*^*2*^ gene and a variety that does not (M82), and only found even coverage along chromosome 9 with the LYC 1969 sample, as expected (Supplementary Fig. 7)(Alonge *et al*., 2020). Further to this, we aligned unassembled short DNA sequences from two other greenhouse tomato cultivars harboring the *Tm-2*^*2*^ gene, Moneymaker-TMV (MmTMV) and Merlice, with SL4.0. We found that the MmTMV introgression appears to be identical to that found in our MbTMV assembly, whereas the Merlice introgression is estimated to be at least 2.67 Mbp longer (Fig 3d & Supplementary Fig. 6).

**Fig. 3:**
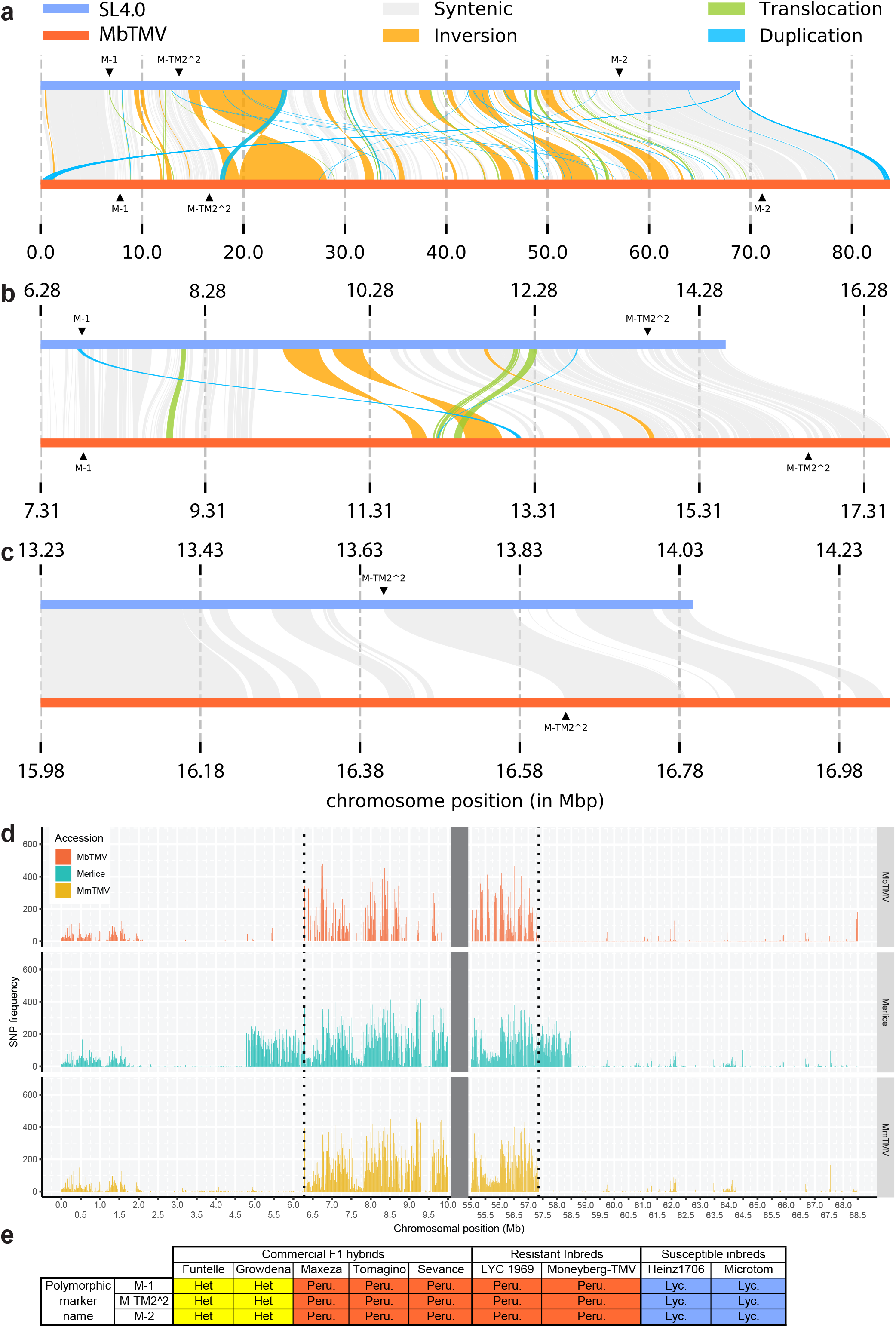
Linkage drag of *S. peruvianum* derived sequence on chromosome 9. **a**, Synteny and Rearrangement (SyRI) plot of MbTMV chromosome 9 (Query) aligned against SL4.0 chromosome 9 (Ref). **b**, SyRI plot of ∼10 Mbp region including TMV resistance gene locus. **c**, SyRI plot of ∼1 Mbp region including TMV resistance gene locus. For **a, b** and **c**, re-alignments were performed on the specific regions and genomic positions (in Mb) are indicated on SL4.0 (Ref) and MbTMV (Query). M-1, M-TM2^2 and M-2 indicate genetic marker positions. Non-syntenic (non-matching) regions are shown as white gaps between Ref and Query. **d**, MbTMV, MmTMV (Moneymaker-TMV) and Merlice variant calling against SL4.0. SNP frequency (calculated in a 10kb window) plotted over the first 10 Mbp and last 13.5 Mbp of SL4.0 chromosome 9. *S. peruvianum* introgression was identified by increased SNP frequency. The two dotted vertical lines indicate introgression start and endpoints for MbTMV and MmTMV. **e**, Marker studies on the TMV resistance introgression from *S. peruvianum* including 5 commercial hybrids: Maxeza (Truss tomato), Tomagino (Cherry tomato), Funtelle (Date tomato), Growdena (Beefsteak tomato) and Sevance (“on the vine” tomato), 2 inbred lines resistant to TMV (LYC1969 and Moneyberg-TMV) and 2 inbred lines susceptible to TMV (Heinz 1706 and Microtom). Marker locations of markers M-1, M-TM2^2 and M-2 are shown in **a**.

The *Tm-2*^*2*^ gene is present in the majority of modern tomato greenhouse cultivars, therefore we set out to check the size of the introgression in five commercial hybrids that were generated in three different breeding programs (Fig. 3e, Supplementary Fig. 8 & Supplementary Fig. 9). To this end we used our assembly to design two polymorphic markers: one at the upstream extreme of the introgression (M-1) and a second at the downstream extreme of the introgression (M-2) (Fig. 3a & Supplementary Fig. 8). Using a published marker within the *Tm-2*^*2*^ gene (M-TM2^2), we found that all five hybrids contained the *Tm-2*^*2*^ gene, two in the heterozygous state (both produced by Syngenta), whereas the other three were homozygous (Fig. 3e & Supplementary Fig. 8). The two hybrids heterozygous for the M-TM2^2 marker were also heterozygous at M-1 and M-2, while the other three hybrids were homozygous at M-1 and M-2 (Fig. 3e & Supplementary Fig. 8). The absolute genetic linkage of the three markers in three different, and heavily resourced, tomato breeding programs confirms that conventional backcross breeding has been unable to break down the introgression.

## Discussion

Here, we have assembled a tomato genome *de novo* to chromosome level contigs using long read DNA sequences alone. The raw HiFi and ONT assemblies were highly contiguous and had different breakpoints in the respective assemblies, therefore merging the assemblies led to further increases in contiguity. Our merged assembly compares favourably (using BUSCO, LAI and chromosome length metrics) with previous tomato genome assemblies that relied on reference- or HiC-based scaffolding (Bolger *et al*., 2014; Schmidt *et al*., 2017; Hosmani *et al*., 2019; Alonge *et al*., 2019, 2020; Wang *et al*., 2020*b*). We made use of the merged assembly to explore the fully assembled 64.1 Mbp introgression from *S. peruvianum* where the *Tm-2*^*2*^ gene, conferring resistance to ToMV and TMV, resides.

Multiple technical developments have facilitated this highly contiguous tomato genome assembly. Firstly, HiFi reads are highly accurate and downsampling revealed that for inbred tomato as low as 20.3 Gbp (equal to about 24x coverage of our final merged assembly) was sufficient for an assembly with an N50 of 20.8 Mbp, with a largest contig of 66.3 Mbp (Supplementary Table 3). Secondly, HMW DNA extraction coupled with Circulomics long read enrichment can now reliably lead to ONT PromethION yields of at least 80 Gbp of usable data, with mean read lengths of at least 20 kbp. The longer ONT reads can facilitate assemblies with more complete chromosomes (Supplementary Fig. 1), yet gene completeness was slightly lower likely due to higher remaining uncorrectable base errors (Table 2). Thirdly, both HiFi and ONT assembly tools are now computationally efficient and allow the testing of various parameters and input data sets. Our HiFi data was assembled by Hifiasm in less than 2 hours with 128 threads, while Flye offers a fast but not as contiguous choice for the ONT platform. Even when using the NECAT assembler, it only takes 3 days with 256 threads to assemble a tomato genome. Finally, by merging the HiFi and ONT assemblies, which had different breakpoints, we were able to harness the complementarity of the two technologies. A recent comparison pointed out that PacBio HiFi reads tend to lead to better assembly of the barley genome than ONT (Mascher *et al*., 2021), while here we showed that the two platforms can be complementary. Highly accurate HiFi reads, and the longer, less accurate, ONT reads facilitated an assembly that spans many complex repeated sequences. As in recent human, Arabidopsis and banana genome assemblies (Belser *et al*., 2021; Nurk *et al*., 2021; Naish *et al*., 2021), the 5S rDNA and 45s rDNA sequences were not complete in our assembly, while some tomato centromeres and telomeres appear to be complete, including a convincing telomere-to-telomere (Hifiasm) assembly of chromosome 5 (Supplementary Fig. 4). Both technologies yielded assemblies that would have been impossible just two years ago, and it appears likely that higher quality ONT long read data will further improve assembly metrics in the near future. The approach we have used here may be used as a simple recipe in other similar sized, homozygous, genomes.

The *Tm-2*^*2*^ gene was known to be within a large introgression (Lin *et al*., 2014) but the complete structure of the region has until now remained unassembled. Despite almost six decades of breeding, and passing through likely hundreds of backcrosses, there remains absolute linkage between the *Tm-2*^*2*^ gene and the two extremes of the introgression - a classic case of linkage drag (Fig. 3e) (Pelham, 1966; Lin *et al*., 2014; Schouten *et al*., 2019). The introgression in Moneyberg-TMV and Moneymaker-TMV is exactly the same length, agreeing with primary accounts of introgression history (see Materials and Methods), while the commercial hybrid Merlice, surprisingly, has a slightly longer introgression (Fig. 3d). The introgression contains several large inversions but the *Tm-2*^*2*^ gene itself is found in a region syntenic to *S. lycopersicum* (Fig. 3 a-c). Pericentromeric heterochromatin makes up about 70% of the tomato genome and meiotic crossovers rarely occur within these regions (Lhuissier *et al*., 2007; Demirci *et al*., 2017; De Haas *et al*., 2017). The *Tm-2*^*2*^ gene is located within the pericentromeric heterochromatin and that region does not recombine in a Moneymaker x *S. pimpinellifolium* RIL population (where no *Tm-2*^*2*^ gene introgression is present), suggesting that the *Tm-2*^*2*^ gene position on chromosome 9 pre-disposes it to linkage drag. However, in the same RIL population crossovers do occur up to ∼1.9 Mbp proximal of the start of the introgression (De Haas *et al*., 2017). This suggests that the higher polymorphism rate and structural variation between *S. lycopersicum* and *S. peruvianum* (compared to the more closely related *S. lycopersicum* and *S. pimpinellifolium)* likely partially explain the extreme linkage drag in this region.

Why do meiotic crossovers not occur within regions of the introgression that are syntenic to *S. lycopersicum*? It remains to be determined whether meiotic DNA double strand breaks (DSBs), a prerequisite for crossovers, occur within the introgression and whether in hybrids that are hemizygous for the introgression chromosome 9 can completely synapse. Given that the introgression contains many genes, and gene promoters/terminators are targets for meiotic DSBs in plants (Choi *et al*., 2018) it is probable that meiotic DSBs do occur within the introgression region, even in hemizygous hybrids. DSBs inside the introgression are likely repaired as non-crossovers. It is possible that the modulation of factors that can change chromosomal distributions of meiotic crossover (e.g. non-CG DNA methylation) could open up this region to meiotic recombination (Underwood *et al*., 2018; Zhao *et al*., 2021; Wang *et al*., 2021).

Introgression of resistance genes from wild crop relatives can have negative side effects, such as reduced yield and quality, due to linkage drag (Tanksley *et al*., 1998; Rubio *et al*., 2016; Chitwood-Brown *et al*., 2021). Employing the *Tm-2*^*2*^ gene introgression in a hemizygous state leads to a higher yield in field tomatoes (Tanksley *et al*., 1998). However, only two of the six commercial hybrids we studied keep the introgression in a hemizygous state (Fig. 3e) (Schouten *et al*., 2019), which may be due to smaller yield gains of a hemizygous introgression in greenhouse varieties or to reduce the risk of ToMV/TMV infection during the production of hybrid seeds. Nonetheless, the *Tm-2*^*2*^ gene introgression from *S. peruvianum* alters the levels of more than three hundred metabolites (Zhu *et al*., 2018), suggesting factors linked to the *Tm-2*^*2*^ gene compromise multiple important tomato traits. We expect that the rapid generation of near complete genome assemblies will be exploited in future to decode additional introgressions from wild crop relatives, and subsequent genome engineering will lead to resistant crop varieties with better yield and taste.

## Materials and Methods

### Plant materials and growth

Moneyberg-TMV was originally developed at De Ruiter seeds (Bleiswijk, South Holland, Netherlands) by introgression of the *Tm-2*^*2*^ gene from a source that contained *S. peruvianum* germplasm originally provided by Alexander (Alexander 1963). The *S. lycopersium* basis of the line originates from the Moneyberg cultivar, an open pollinated, indeterminate, greenhouse tomato cultivar, that was developed by selection from Moneymaker at Vandenberg seeds (Gebr. van den Berg, Naaldwijk, South Holland, Netherlands). Moneymaker itself was first developed in the early 1900s by Fred Stonor (Southampton, England, UK). Maxeza F1 (Truss tomato) and Tomagino F1 (Cherry tomato) were produced by Enza Zaden. Funtelle F1 (Date tomato) and Growdena F1 (Beefsteak tomato) were produced by Syngenta. Sevance F1 (“on the vine” tomato) and Merlice F1 (Truss tomato) were produced by De Ruiter seeds. Moneymaker-TMV seeds were from the stocks of WUR-Plant breeding which were originally provided by De Ruiter seeds. For further seed origin information please see the acknowledgements.

For genome sequencing and meiotic cytology, ten Moneyberg-TMV plants (all progeny of a single highly-inbred parent) were grown in the MPIPZ greenhouses during the late spring of 2020 under natural light supplemented with artificial light to ensure 16 hours light per day. Young unexpanded leaves were collected from 5-week-old plants and snap frozen in liquid nitrogen before interim storage at -80 degrees. For whole-genome resequencing, Moneymaker-TMV and Merlice plants were grown in the greenhouses of WUR until the third full leaf, and shoot tips with young leaves were harvested and snap frozen in liquid nitrogen. For marker studies, Moneyberg-TMV, LYC1969, Microtom, Heinz1706, Maxeza, Tomagino, Funtelle, Growdena and Sevance seeds were germinated *in vitro* on 0.8% agarose.

### Chromosome spreading

Appropriate stages (3-4mm) of meiotic buds from Moneyberg-TMV were fixed in 1mL fresh Carnoy’s solution composed of 3:1 (v/v) absolute ethanol:glacial acetic acid, and placed under vacuum for at least 30 minutes to ensure tissue infiltration. The Carnoy’s solution was replaced and the samples kept at room temperature for 48 hours until they turned white. The fixed materials were used for chromosome spreading essentially as previously described (Ross *et al*., 1996). In brief, 50-60 meiotic anthers were isolated from meiotic buds and placed into a 50 µL enzyme mixture (0.3% cellulase, 0.3% cytohelicase and 0.3% pectolyase in citric buffer 10 mM, pH 4.6) to digest for 2.5 hours at 37 °C. 2-3 anthers were used to make a single slide. After drying 8 µL DAPI (2 µg/mL) was added to stain the chromosomes. Finally, meiocytes images were taken using a Zeiss Axio Imager Z2 Microscope and images were analysed using the Zeiss ZEN 2 (blue edition) software.

### Moneyberg-TMV High molecular weight DNA extraction

High molecular weight (HMW) DNA of Moneyberg-TMV was isolated from 1.5 grams of young leaf material with a NucleoBond HMW DNA kit (Macherey Nagel). DNA quality was assessed with a FEMTOpulse device (Agilent) and quantity measured using a Quantus Fluorometer (Promega).

### Moneyberg-TMV Library preparations and sequencing

A HiFi library was prepared according to the manual “Procedure & Checklist - Preparing HiFi SMRTbell® Libraries using SMRTbell Express Template Prep Kit 2.0” with an initial DNA fragmentation by g-Tubes (Covaris) and final library size binning into defined fractions by SageELF (Sage Science). Size distribution was again controlled by FEMTOpulse (Agilent). Size-selected libraries with an expected insert size of 15 kbp and 18 kbp were independently sequenced on single SMRT cells of a Pacific Biosciences Sequel II device at the MPGC Cologne with Binding kit 2.0 and Sequel II Sequencing Kit 2.0 for 30 h (Pacific Biosciences).

9 µg of HMW DNA was size selected using the Circulomics Short-Read Eliminator XL Kit (Circulomics Cat# SKU SS-100-111-01). Out of this size selected DNA 1.5 µg was used as input for two library preparations with the Oxford Nanopore LSK-110 ligation sequencing Kit. Half of each library was loaded onto a PromethION PRO-002 Flowcell (R9.4.1 pores) and sequenced for 24 h. Then the Flowcells were rinsed using the Oxford Nanopore Wash Kit WSH-003 and the second part of each library was loaded and sequencing continued for another 48 h. ONT Sequencing was performed on an Oxford Nanopore P24 PromethION at the Forschungszentrum Jülich using MinKNOW version 21.02.7 without live basecalling. Basecalling of the Oxford Nanopore read data was done using guppy basecaller version 5.0.11 using the R9.4.1 PromethION super high-accuracy model (dna_r9.4.1_450bps_sup_prom) including filtering out basecalled reads with an average Phred-Score of less than 9.0. Reads with a Q-value below 9 were excluded before further analysis, including genome assembly. To remove adapter sequences and split chimeric reads the remaining Nanopore reads were then filtered further with porechop version 0.2.4 with default settings.

A chromatin conformation capture library was prepared using 0.5 grams of young leaf material as the input. All treatments were according to the recommendations of the kit vendor (Omni-C, Dovetail) for plants. As a final step, an Illumina-compatible library was prepared (Dovetail) with an insert size of 540bp and sequenced (paired-end 2 × 150 bp) on an Illumina HiSeq 3000 device at the MPGC Cologne to generate 11.37 million PE reads.

### Whole genome re-sequencing

Moneymaker-TMV and Merlice plant material was ground into fine powder by mortar and pestle, and DNA was isolated by a CTAB based method (Healey *et al*., 2014). DNA was dissolved in ultrapure water and DNA was checked by gel, Qubit and nanodrop. Library preparation and whole genome sequencing was performed by Novogene using the Illumina HiSeq platform.

### Genome assemblies and assembly statistics

Hifiasm v0.14.2 (Cheng *et al*., 2021) used to assemble the HiFi reads with default settings. Canu v2.1.1 was used to assemble the HiFi reads in “HiCanu” mode (Nurk *et al*., 2020) with an estimated genome size of 916 Mbp based on kmer counting of the raw HiFi data using Jellyfish v2.2.6 (Marçais and Kingsford, 2011).

Flye v2.8.2 (Kolmogorov *et al*., 2019) was used to assemble the ONT reads with the option --nano-raw for unpolished Nanopore reads. NECAT version 0.0.1_update20200803 was used to assemble the ONT reads (Chen *et al*., 2021) with default options apart from increasing the coverage used for assembly from 30x to 40x by setting PREP_OUTPUT_COVERAGE to 40.

ONT and HiFi reads were randomly subsampled in 9 sets of 90, 80, 70, 60, 50, 40, 30, 20 and 10% of the original reads and used for assemblies using NECAT (ONT) and Hifiasm (HiFi) with the same settings as for the original assemblies.

Quast v5.0.2 (Mikheenko *et al*., 2018) and GAAS v1.1.0 (https://github.com/NBISweden/GAAS) were used to calculate statistics on fasta files. Benchmarking Universal Single Copy Orthologs (BUSCO) was calculated using BUSCO v5.2.1 (Seppey *et al*., 2019) depending on hmmsearch v3.1 and metaeuk v4.a0f584d, was used with lineage datasets solanales (https://busco-data.ezlab.org/v5/data/lineages/solanales_odb10.2020-08-05.tar.gz) and eudicots (https://busco-data.ezlab.org/v5/data/lineages/eudicots_odb10.2020-09-10.tar.gz) to obtain evolutionarily-informed expectations of gene content. To assess the LTR assembly index (LAI) LTR retriever v2.9.0 (Ou *et al*., 2018) was run with default settings on the respective genome assemblies.

### Scaffolding approaches

The NECAT assembly was polished with all of the ONT reads using Medaka v1.2.3 (https://nanoporetech.github.io/medaka/) and the model specific for PromethION R.9.4.1 data basecalled with guppy 5.0.11 in “super high accuracy” mode. The Flye assembly was also polished as described above.

Next, the NECAT 40x medaka and Flye v2.8.2 medaka assemblies were merged. This was done by running nucmer (part of mummer v.4.0.0rc1) with the -l parameter to prevent invalid contig links using the NECAT assembly as query and the Flye assembly as reference. The alignment table was then filtered with delta-filter (also part of mummer) using the options -r -q -i 0.95 to only use reciprocal best matches per region and a minimum identity of 95% in overlaps. This matches the default settings in the quickmerge wrapper which still relies on mummer version 3. Finally quickmerge (Chakraborty *et al*., 2016) was used with the parameter -c 7.0 to increase stringency of merging regions to prevent assembly artifacts. The other parameters mandatory for quickmerge were set to the same values as used in the wrapper script by default (-hco 5.0 -l 0 -ml 5000). The resulting merged assembly was subsequently polished with all HiFi reads using Racon v1.4.3 (Vaser *et al*., 2017) together with Minimap version 2.21 (Li, 2018) using the “map-hifi” preset for mapping the reads to the genome to overcome uncorrectable errors resulting from Nanopore reads.

In parallel, the Canu and Hifiasm assemblies were merged using the Canu assembly as reference and the Hifiasm assembly as query with quickmerge (as above) but using the default -c parameter of 1.5. Due to the low baseline error rate of HiFi Reads no further polishing was done on these assemblies.

The resulting merged (and polished) assemblies were then merged again using the Nanopore (Flye+NECAT) as query and the PacBio (Hifiasm+Canu) as reference using the same default parameters as for merging the two PacBio assemblies before. The final merged assembly was named MbTMV.

All assemblies were mapped to SL4.0 (Hosmani *et al*., 2019) using Minimap2 v2.17 (Li, 2018) with default settings, followed by scaffolding using RaGOO v1.11 (Alonge *et al*., 2019) with default options.

### Dot plots

D-GENIES version 1.2.0 (Cabanettes and Klopp, 2018) was run with default settings with alignments generated using Minimap2 v2.21 (Li, 2018) providing the *Solanum lycopersicum* cv. ‘Heinz1706’ SL4.0 (Hosmani *et al*., 2019) as reference (target) and our assembly as query. For legibility the chromosome names in both assemblies were shortened to “ch00 - ch12”. Local installations of reDOTable v1.1 (https://www.bioinformatics.babraham.ac.uk/projects/redotable/ or github https://github.com/s-andrews/redotable) were used to generate dotplots.

### Validation of MbTMV assembly by Omni-C

Following the esrice/hic-pipeline (https://github.com/esrice/hic-pipeline), Omni-C (Dovetail) paired end reads were mapped separately using Burrows-Wheeler Aligner v0.7 (Li and Durbin, 2009) and the filter_chimeras.py script. Mapped reads were combined using the combine_ends.py script, with 3 iteration steps and a minimum mapping quality value of 20 (default), followed by adding read mate scores, sorting and removing duplicate reads using Samtools v1.9 (Danecek *et al*., 2021). The resulting bam file was converted to a bed file and sorted (-k 4) using Bedtools v2.30 (Quinlan and Hall, 2010). Scaffolding was done using Salsa v2.2 (Ghurye *et al*., 2019) with optional settings: -e DNASE -m yes -p yes, until no more breakpoints were visualized.

A modified version of the convert.sh script was used to convert Salsa2 produced scaffolds_FINAL.fasta and scaffolds_FINAL.agp to Hi-C file, which was used within Juicebox (https://github.com/aidenlab/Juicebox) v1.11.08 to generate a Hi-C contact plot.

### Validation of MbTMV assembly by re-mapping raw data and circos plot

ONT data was mapped using Minimap2 version 2.18-r1028-dirty (Li, 2018), with the preset “map-ont” whereas PacBio HiFi data was mapped with the preset “map-hifi”. The output SAM file was converted, sorted and indexed using Samtools v1.9 (Danecek *et al*., 2021). Samtools v1.9 (Danecek *et al*., 2021) was used to extract the coverage per base, including positions with zero depth, from the bam file. The resulting average depth was summarized in overlapping windows of 200 kbp every 100 kbp and visualized using NG-circos (Cui *et al*., 2020). In addition, sequences of the 45S rDNA intergenic spacer, 5S rDNA, 45S rDNA, telomeric repeat and TGR1-4 were searched in MbTMV using blastn and counted in non-overlapping windows of size 1 Mbp.

### Genome alignment, SV detection and further genome analysis

Genomes were aligned to SL4.0 (Hosmani *et al*., 2019), using Minimap2 v2.17 (Li, 2018) with options -ax asm5 --eqx, followed by running SyRI v1.4 (Goel *et al*., 2019), to identify and call polymorphisms and structural variations, with optional settings: -k -F S --log WARN. Plotsr v5 (Goel *et al*., 2019) was used with options -B <annotation.bed> -H <1-9> and -W <1-9> to generate SyRI plots. Fasta headers of genomes were manually formatted to match between genomes, using sed command line, and chromosome 0 was removed, using awk command line, before aligning the genomes. Chromosome 1, 2, 7, 8 and 12 of the MbTMV assembly were reverse complemented before alignment, using cat command line.

Using the SyRI generated vcf output of the whole chromosome alignment (Supplementary Table 6), sequences used for Figures 3B and 3C were extracted (SL4.0:6286825-14549782 MbTMV:7314513-17630288 and SL4.0:13227892-14039189 MbTMV:15980168-17044082 for figure 3B and 3C, respectively), using Samtools v1.9 (Danecek *et al*., 2021). Sequences were aligned using Minimap2 v2.17 (Li, 2018) with options -ax asm5 --eqx followed by running SyRI v1.4 and Plotsr v5 (Goel *et al*., 2019). Figures 3B and 3C were manually edited using Adobe Illustrator 25.2.3 adding genomic positions based on extracted regions.

Chromosome lengths were extracted from the fasta files, MbTMV, *S. lycopersicum* cv. ‘Heinz1706’ SL4.0 (Hosmani *et al*., 2019), obtained from ftp://ftp.solgenomics.net/genomes/Solanum_lycopersicum/Heinz1706/assembly/build_4.00/S_lycopersicum_chromosomes.4.00.fa, *S. pennellii* ‘LA0716’ (Bolger *et al*., 2014), obtained from ftp://ftp.solgenomics.net/genomes/Solanum_pennellii/Spenn.fasta, *S. pimpinellifolium* ‘LA2093’ v1.5 (Wang *et al*., 2020*b*) obtained from ftp://ftp.solgenomics.net/genomes/Solanum_pimpinellifolium/LA2093/Spimp_LA2093_genome_v1.5/LA2093_genome_v1.5.fa, using awk command line and plotted within R version 4.0.3 (2020-10-10) (https://cran.r-project.org/) using ggplot2 and the tidyverse package (https://www.tidyverse.org/).

### Further analysis of chromosome 9 introgression

M82 and LYC 1969 ONT data (Alonge *et al*., 2020) were mapped to both SL4.0 and MbTMV using Minimap2 v2.17 (Li, 2018), with the options -ax map-ont --eqx. The output SAM file was converted, sorted and indexed using Samtools v1.9 (Danecek *et al*., 2021). Coverage plots per chromosome were constructed with goleft IndexCov v0.2.4 (BS *et al*., 2017).

Moneymaker and Merlice illumina data was mapped to SL4.0 using bowtie2 v2.2.8 with options -q --phred33 --very-sensitive --no-unal -p 12 -x <SL4.0.fasta> -U reads1.fq,reads2.fq -S. Variant calling was done using bcftools v1.9 mpileup and call with options -mv -Ov (Li, 2011) (https://samtools.github.io/bcftools/bcftools.html).

SNP density was extracted from MbTMV (SyRi output), MoneymakerTMV and Merlice vcf files using VCFtools v.0.1.16 (Danecek *et al*., 2011). Plots were made within R 4.0.3 (2020-10-10) (https://cran.r-project.org/) using ggplot2 and the tidyverse package (https://www.tidyverse.org/).

### Marker studies

Cotyledons from *in vitro* germinated samples were collected and lyophilized overnight in an Alpha 1-4 freeze dryer (Martin Christ GmbH). DNA extraction was performed using the BioSprint 96 DNA Plant kit protocol (Qiagen) and eluted in 200µl AE buffer. M-1 was amplified using the primers dCUn0256 (GGAACCTCGAGTCCTTACAGT) and dCUn0257 (GGCAGCCATTAGCAAACCCA). M-TM2^2 was amplified using published primers (Lanfermeijer *et al*., 2005). M-2 was amplified using the primers dCUn0258 (TTCTGTTCCAGACCCGACCT) and dCUn0259 (CCCATTAACCTCCAGACGGG). 35 rounds of PCR were performed using Mango Taq (Bioline) under standard conditions. M-1 was digested with AflII (NEB) overnight at 37°C, M-TM2^2 was digested with BslI (NEB) for 3 hours at 55°C and M-2 was digested with XhoI (NEB) overnight at 37°C. 10µl per restriction digest was loaded on a 3% agarose gel.

## Data availability

The assemblies presented in this pre-print are made available at https://www.plabipd.de/portal/moneyberg

## Acknowledgements

Moneyberg-TMV seeds were kindly provided for research purposes by Cilia Lelivelt and Maarten Verlaan (Rijk Zwaan, Netherlands). Maxeza F_1_ and Tomagino F_1_ tomato seeds were kindly provided for research purposes by Martijn van Stee (Enza Zaden, Netherlands). Funtelle F_1_ and Growdena F_1_ tomato seeds were kindly provided for research purposes by Axel Voss (Syngenta, Germany). Sevance F_1_ seeds (De Ruiter seeds, part of Bayer Group) were purchased for research purposes from Volmary GmbH (Münster, Germany). Merlice F_1_ seeds (De Ruiter seeds, part of Bayer Group) were provided by Tomato World (Netherlands) to HJS for research purposes. Microtom seeds were originally sourced from the Tomato Growers Supply company (Ft. Myers, Florida, USA). LYC1969 seeds were a gift from Zachary Lippman (CSHL, USA). Heinz 1706 seeds were provided by the Centre for Genetic Resources (Wageningen, Netherlands). We are grateful to Gerard Bijsterbosch (WUR) for isolating DNA from Moneymaker-TMV and Merlice. We thank Saurabh Pophaly (MPIPZ), Manish Goel (MPIPZ) and Danny Esselink (WUR) for assistance with bioinformatic analysis. We thank Korbinian Schneeberger (MPIPZ), Quentin Gouil (Walter and Eliza Hall Institute) and Alex Canto-Pastor (UC Davis) for comments on the manuscript. Information on the breeding history of Moneymaker, Moneyberg and Moneyberg-TMV varieties was kindly provided by Theo Schotte (Solynta, Netherlands) and Maarten Koornneef (WUR/MPIPZ). This work was supported by the Max Planck Society, a PhD fellowship awarded to M. W. A. M. Z. from the Malaysian Agricultural Research and Development Institute (MARDI), grants awarded to B. U. (DFG US 98/23, CEPLAS EXC2048 and BMBF 031A536C) and the WUR Department of Plant Breeding (H.S.).

## References

Aflitos S, Schijlen E, de Jong H, et al. 2014. Exploring genetic variation in the tomato (Solanum section Lycopersicon) clade by whole-genome sequencing. The Plant Journal 80, 136–148.

Alexander LJ. 1963. Transfer of a dominant type of resistance to the four known Ohio pathogenic strains of tobacco mosaic virus (TMV) from Lycopersicon peruvianum to L. esculentum. Phytopathology 53, 896.

Alonge M, Soyk S, Ramakrishnan S, Wang X, Goodwin S, Sedlazeck FJ, Lippman ZB, Schatz MC. 2019. RaGOO: fast and accurate reference-guided scaffolding of draft genomes. Genome Biology 2019 20:1 20, 1–17.

Alonge M, Wang X, Benoit M, et al. 2020. Major Impacts of Widespread Structural Variation on Gene Expression and Crop Improvement in Tomato. Cell 182, 145-161.e23.

Bai Y, Lindhout P. 2007. Domestication and breeding of tomatoes: What have we gained and what can we gain in the future? Annals of Botany 100, 1085–1094.

Belser C, Baurens F-C, Noel B, et al. 2021. Telomere-to-telomere gapless chromosomes of banana using nanopore sequencing. bioRxiv, 2021.04.16.440017.

Berlin K, Koren S, Chin CS, Drake JP, Landolin JM, Phillippy AM. 2015. Assembling large genomes with single-molecule sequencing and locality-sensitive hashing. Nature Biotechnology 33, 623–630.

Bolger A, Scossa F, Bolger ME, et al. 2014. The genome of the stress-tolerant wild tomato species Solanum pennellii. Nature Genetics 46, 1034–1038.

BS P, RL C, ME T, AR Q. 2017. Indexcov: fast coverage quality control for whole-genome sequencing. GigaScience 6.

Cabanettes F, Klopp C. 2018. D-GENIES: dot plot large genomes in an interactive, efficient and simple way. PeerJ 6, e4958.

Chakraborty M, Baldwin-Brown JG, Long AD, Emerson JJ. 2016. Contiguous and accurate de novo assembly of metazoan genomes with modest long read coverage. Nucleic Acids Research 44, e147.

Chang S Bin, Yang TJ, Datema E, et al. 2008. FISH mapping and molecular organization of the major repetitive sequences of tomato. Chromosome Research 16, 919–933.

Chen Y, Nie F, Xie S-Q, et al. 2021. Efficient assembly of nanopore reads via highly accurate and intact error correction. Nature Communications 2021 12:1 12, 1–10.

Cheng H, Concepcion GT, Feng X, Zhang H, Li H. 2021. Haplotype-resolved de novo assembly using phased assembly graphs with hifiasm. Nature Methods 2021 18:2 18, 170–175.

Chitwood-Brown J, Vallad GE, Lee TG, Hutton SF. 2021. Characterization and elimination of linkage-drag associated with Fusarium wilt race 3 resistance genes. Theoretical and Applied Genetics 1, 3.

Choi K, Zhao X, Tock AJ, et al. 2018. Nucleosomes and DNA methylation shape meiotic DSB frequency in Arabidopsis thaliana transposons and gene regulatory regions. Genome research 28, 532–546.

Cui Y, Cui Z, Xu J, Hao D, Shi J, Wang D, Xiao H, Duan X, Chen R, Li W. 2020. NG-Circos: next-generation Circos for data visualization and interpretation. NAR Genomics and Bioinformatics 2.

Danecek P, Auton A, Abecasis G, et al. 2011. The variant call format and VCFtools. Bioinformatics 27, 2156–2158.

Danecek P, Bonfield JK, Liddle J, et al. 2021. Twelve years of SAMtools and BCFtools. GigaScience 10, 1–4.

Demirci S, van Dijk ADJ, Sanchez Perez G, Aflitos SA, de Ridder D, Peters SA. 2017. Distribution, position and genomic characteristics of crossovers in tomato recombinant inbred lines derived from an interspecific cross between Solanum lycopersicum and Solanum pimpinellifolium. The Plant Journal 89, 554–564.

Dumschott K, Schmidt MHW, Chawla HS, Snowdon R, Usadel B. 2020. Oxford Nanopore sequencing: new opportunities for plant genomics? Journal of Experimental Botany 71, 5313–5322.

Foolad MR, Stoltz T, Dervinis C, Rodriguez RL, Jones RA. 1997. Mapping QTLs conferring salt tolerance during germination in tomato by selective genotyping. Molecular Breeding 3, 269–277.

Ganal MW, Lapitan NL, Tanksley SD. 1991. Macrostructure of the tomato telomeres. The Plant Cell 3, 87–94.

Gao L, Gonda I, Sun H, et al. 2019. The tomato pan-genome uncovers new genes and a rare allele regulating fruit flavor. Nature Genetics 51, 1044–1051.

Ghurye J, Rhie A, Walenz BP, Schmitt A, Selvaraj S, Pop M, Phillippy AM, Koren S. 2019. Integrating Hi-C links with assembly graphs for chromosome-scale assembly. PLOS Computational Biology 15, e1007273.

Goel M, Sun H, Jiao WB, Schneeberger K. 2019. SyRI: finding genomic rearrangements and local sequence differences from whole-genome assemblies. Genome Biology 20, 1–13.

De Haas LS, Koopmans R, Lelivelt CLC, Ursem R, Dirks R, James GV. 2017. Low-coverage resequencing detectsmeiotic recombination pattern and features in tomato RILs. DNA Research 24, 549–558.

Healey A, Furtado A, Cooper T, Henry RJ. 2014. Protocol: a simple method for extracting next-generation sequencing quality genomic DNA from recalcitrant plant species. Plant Methods 2014 10:1 10, 1–8.

Hon T, Mars K, Young G, et al. 2020. Highly accurate long-read HiFi sequencing data for five complex genomes. Scientific Data 7, 1–11.

Hosmani P, Flores-Gonzalez M, van de Geest H, et al. 2019. An improved de novo assembly and annotation of the tomato reference genome using single-molecule sequencing, Hi-C proximity ligation and optical maps. bioRxiv, 767764.

Jo S-H, Koo D-H, Kim JF, Hur C-G, Lee S, Yang T, Kwon S-Y, Choi D. 2009. Evolution of ribosomal DNA-derived satellite repeat in tomato genome. BMC Plant Biology 9, 42.

Kim HT, Lee JM. 2018. Organellar genome analysis reveals endosymbiotic gene transfers in tomato. PLoS ONE 13.

Kolmogorov M, Yuan J, Lin Y, Pevzner PA. 2019. Assembly of long, error-prone reads using repeat graphs. Nature Biotechnology 37, 540–546.

Koren S, Schatz MC, Walenz BP, et al. 2012. Hybrid error correction and de novo assembly of single-molecule sequencing reads. Nature Biotechnology 30, 693–700.

Lanfermeijer FC, Dijkhuis J, Sturre MJG, De Haan P, Hille J. 2003. Cloning and characterization of the durable tomato mosaic virus resistance gene Tm-22 from Lycopersicon esculentum. Plant Molecular Biology 52, 1037–1049.

Lanfermeijer FC, Warmink J, Hille J. 2005. The products of the broken Tm-2 and the durable Tm-22 resistance genes from tomato differ in four amino acids. Journal of Experimental Botany 56, 2925–2933.

Lhuissier FGP, Offenberg HH, Wittich PE, Vischer NOE, Heyting C. 2007. The mismatch repair protein MLH1 marks a subset of strongly interfering crossovers in tomato. Plant Cell 19, 862–876.

Li H. 2011. A statistical framework for SNP calling, mutation discovery, association mapping and population genetical parameter estimation from sequencing data. Bioinformatics 27, 2987–2993.

Li H. 2018. Minimap2: Pairwise alignment for nucleotide sequences. Bioinformatics 34, 3094–3100.

Li H, Durbin R. 2009. Fast and accurate short read alignment with Burrows-Wheeler transform. Bioinformatics 25, 1754–1760.

Li J, Liu L, Bai Y, Zhang P, Finkers R, Du Y, Visser RGF, van Heusden AW. 2010. Seedling salt tolerance in tomato. Euphytica 178, 403–414.

Lin T, Zhu G, Zhang J, et al. 2014. Genomic analyses provide insights into the history of tomato breeding. Nature Genetics 46, 1220–1226.

Liu Y, Du H, Li P, et al. 2020. Pan-Genome of Wild and Cultivated Soybeans. Cell 182, 162-176.e13.

Marçais G, Kingsford C. 2011. A fast, lock-free approach for efficient parallel counting of occurrences of k-mers. Bioinformatics 27, 764–770.

Mascher M, Wicker T, Jenkins J, et al. 2021. Long-read sequence assembly: a technical evaluation in barley. The Plant Cell 33, 1888–1906.

Michael TP, Jupe F, Bemm F, Motley ST, Sandoval JP, Lanz C, Loudet O, Weigel D, Ecker JR. 2018. High contiguity Arabidopsis thaliana genome assembly with a single nanopore flow cell. Nature Communications 9, 1–8.

Mikheenko A, Prjibelski A, Saveliev V, Antipov D, Gurevich A. 2018. Versatile genome assembly evaluation with QUAST-LG. Bioinformatics 34, i142–i150.

Naish M, Alonge M, Wlodzimierz P, et al. 2021. The genetic and epigenetic landscape of the Arabidopsis centromeres. bioRxiv, 2021.05.30.446350.

Nurk S, Koren S, Rhie A, et al. 2021. The complete sequence of a human genome. bioRxiv, 2021.05.26.445798.

Nurk S, Walenz BP, Rhie A, Vollger MR, Logsdon GA, Grothe R, Miga KH, Eichler EE, Phillippy AM, Koren S. 2020. HiCanu: accurate assembly of segmental duplications, satellites, and allelic variants from high-fidelity long reads. Genome Research 30, gr.263566.120.

Ou S, Chen J, Jiang N. 2018. Assessing genome assembly quality using the LTR Assembly Index (LAI). Nucleic acids research 46, e126.

Payne A, Holmes N, Rakyan V, Loose M. 2019. Bulkvis: A graphical viewer for Oxford nanopore bulk FAST5 files. Bioinformatics 35, 2193–2198.

Pelham J. 1966. Resistance in tomato to tobacco mosaic virus. Euphytica 15, 258–267.

Peralta IE, Spooner DM, Knapp S. 2008. Taxonomy of wild tomatoes and their relatives (Solanum sect. Lycopersicoides, sect. Juglandifolia, sect. Lycopersicon ; Solanaceae). American Society of Plant Taxonomists.

Perry KL, Palukaitis P. 1990. Transcription of tomato ribosomal DNA and the organization of the intergenic spacer. MGG Molecular & General Genetics 221, 102–112.

Powell AF, Courtney LE, Schmidt MH-W, et al. 2020. A Solanum lycopersicoides reference genome facilitates biological discovery in tomato. bioRxiv, 2020.04.16.039636.

Qin P, Lu H, Du H, et al. 2021. Pan-genome analysis of 33 genetically diverse rice accessions reveals hidden genomic variations. Cell 184, 3542-3558.e16.

Quinlan A, Hall I. 2010. BEDTools: a flexible suite of utilities for comparing genomic features. Bioinformatics (Oxford, England) 26, 841–842.

Ross KJ, Fransz P, Jones GH. 1996. A light microscopic atlas of meiosis in Arabidopsis thaliana. Chromosome Research 4, 507–516.

Rubio F, Alonso A, García-Martínez S, Ruiz JJ. 2016. Introgression of virus-resistance genes into traditional Spanish tomato cultivars (Solanum lycopersicum L.): Effects on yield and quality. Scientia Horticulturae 198, 183–190.

Sato S, Tabata S, Hirakawa H, et al. 2012. The tomato genome sequence provides insights into fleshy fruit evolution. Nature 485, 635–641.

Schmidt-Puchta W, Günther I, Sänger HL. 1989. Nucleotide sequence of the intergenic spacer (IGS) of the tomato ribosomal DNA. Plant Molecular Biology 13, 251–253.

Schmidt MHW, Vogel A, Denton AK, et al. 2017. De novo assembly of a new Solanum pennellii accession using nanopore sequencing. Plant Cell 29, 2336–2348.

Schouten HJ, Tikunov Y, Verkerke W, Finkers R, Bovy A, Bai Y, Visser RGF. 2019. Breeding Has Increased the Diversity of Cultivated Tomato in The Netherlands. Frontiers in Plant Science 10, 1606.

Seppey M, Manni M, Zdobnov EM. 2019. BUSCO: Assessing Genome Assembly and Annotation Completeness. Methods in Molecular Biology 1962, 227–245.

Tanksley SD, Bernachi D, Beck-Bunn T, et al. 1998. Yield and quality evaluations on a pair of processing tomato lines nearly isogenic for the Tm2a gene for resistance to the tobacco mosaic virus. Euphytica 99, 77–83.

Underwood CJ, Choi K, Lambing C, et al. 2018. Epigenetic activation of meiotic recombination near Arabidopsis thaliana centromeres via loss of H3K9me2 and non-CG DNA methylation. Genome Research 28, 519–531.

Vaillancourt B, Buell CR. 2019. High molecular weight DNA isolation method from diverse plant species for use with Oxford Nanopore sequencing. bioRxiv, 783159.

Vanburen R, Bryant D, Edger PP, et al. 2015. Single-molecule sequencing of the desiccation-tolerant grass Oropetium thomaeum. Nature 527, 508–511.

Vaser R, Sović I, Nagarajan N, Šikić M. 2017. Fast and accurate de novo genome assembly from long uncorrected reads. Genome Research 27, 737–746.

Vilanova S, Alonso D, Gramazio P, et al. 2020. SILEX: A fast and inexpensive high-quality DNA extraction method suitable for multiple sequencing platforms and recalcitrant plant species. Plant Methods 16, 1–11.

Vollger MR, Logsdon GA, Audano PA, et al. 2020. Improved assembly and variant detection of a haploid human genome using single-molecule, high-fidelity long reads. Annals of Human Genetics 84, 125–140.

Wang J, Chen T, Han M, et al. 2020a. Plant NLR immune receptor Tm-22 activation requires NB-ARC domain-mediated self-association of CC domain. PLoS Pathogens 16, e1008475.

Wang X, Gao L, Jiao C, et al. 2020b. Genome of Solanum pimpinellifolium provides insights into structural variants during tomato breeding. Nature Communications 11.

Wang Y, van Rengs WMJ, Mohd Zaidan MWA, Underwood CJ. 2021. Meiosis in crops: from genes to genomes. Journal of Experimental Botany.

Zapata L, Ding J, Willing EM, et al. 2016. Chromosome-level assembly of Arabidopsis thaliana Ler reveals the extent of translocation and inversion polymorphisms. Proceedings of the National Academy of Sciences of the United States of America 113, E4052–E4060.

Zhao M, Ku JC, Liu B, et al. 2021. The mop1 mutation affects the recombination landscape in maize. Proceedings of the National Academy of Sciences of the United States of America 118, 2009475118.

Zhong XB, Fransz PF, Eden JW Van, Ramanna MS, Van Kammen A, Zabel P, Hans de Jong J. 1998. FISH studies reveal the molecular and chromosomal organization of individual telomere domains in tomato. Plant Journal 13, 507–517.

Zhu G, Wang S, Huang Z, et al. 2018. Rewiring of the Fruit Metabolome in Tomato Breeding. Cell 172, 249-261.e12.

