## Supplementary Figures for "A gap-free tomato genome built from complementary PacBio and Nanopore long DNA sequences reveals extensive linkage drag during breeding"

### Supplementary Figure 1

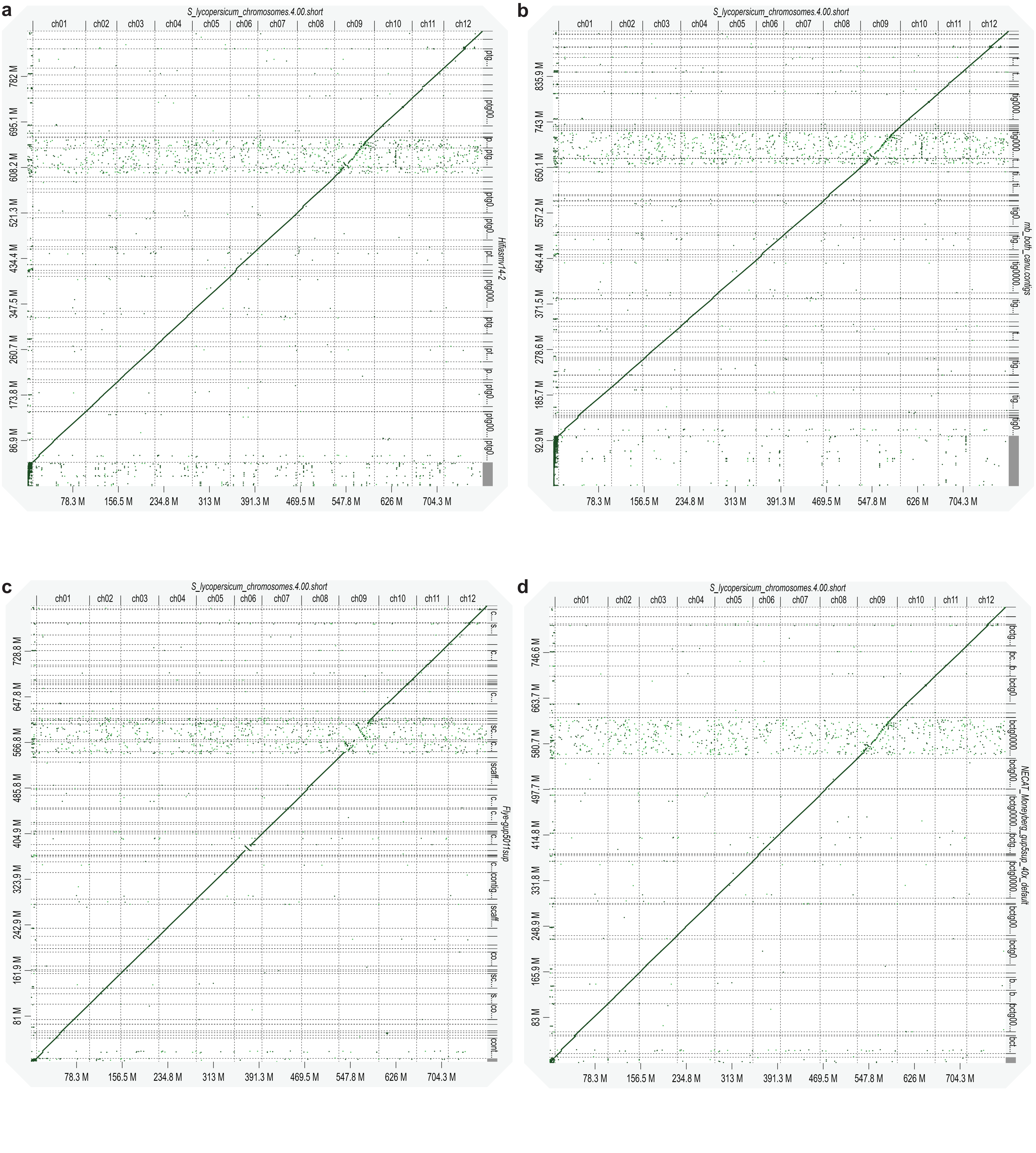


**Supplementary Fig. 1 | Alignment of four Moneyberg-TMV genome assemblies with Heinz 1706 (SL4.0) assembly**

D-GENIES alignments of Hifiasm, Canu, Flye and NECAT assemblies against the SL4.0 assembly. Min identity (abs) was set to 0.65. **a**, SL4.0 (top) versus Hifiasm assembly. **b**, SL4.0 (top) versus Canu assembly. **c**, SL4.0 (top) versus Flye assembly. **d**, SL4.0 (top) versus NECAT assembly

### Supplementary Figure 2

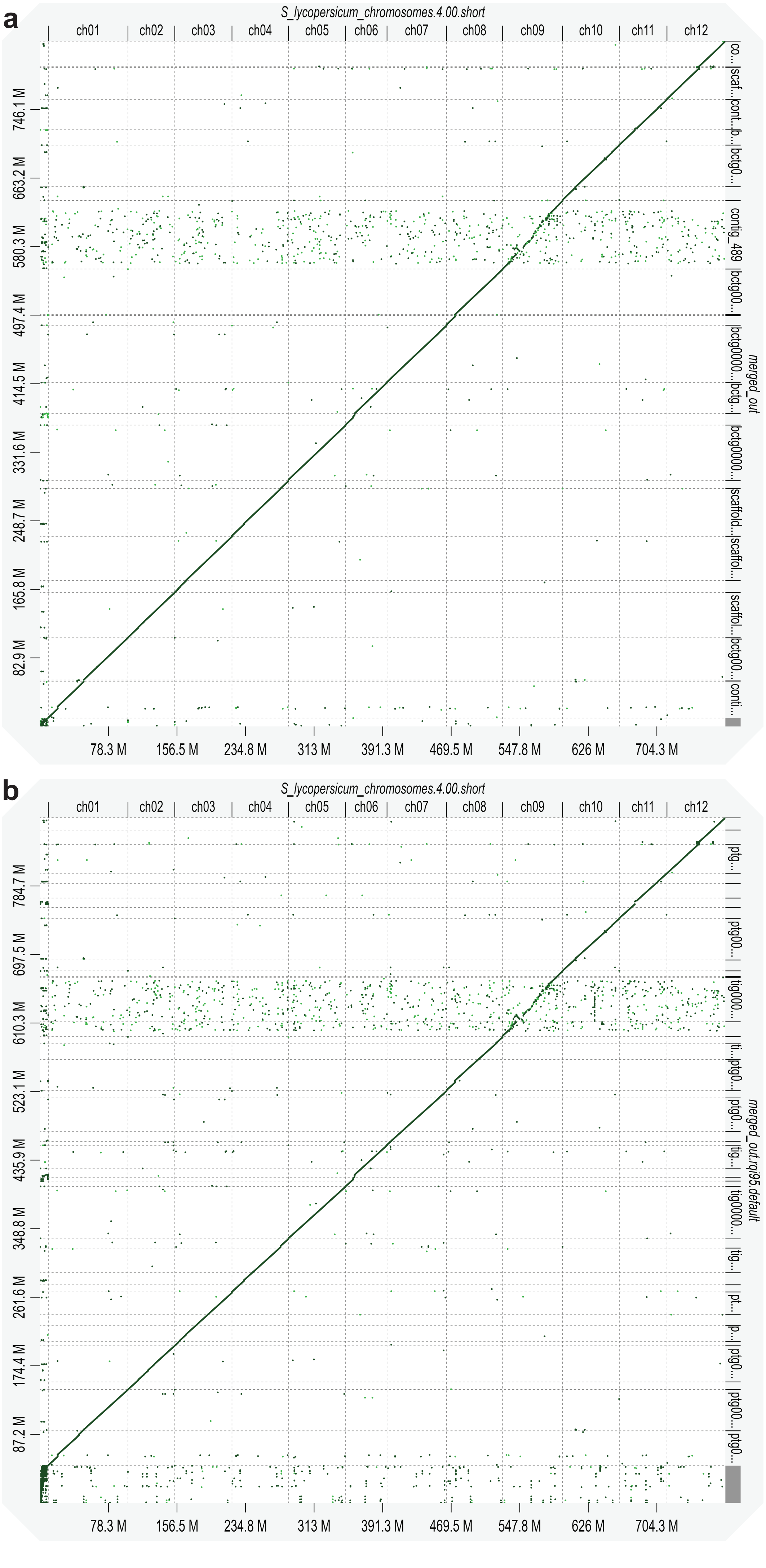


**Supplementary Fig. 2 | Alignment of the merged assemblies to SL4.0**
D-GENIES alignments of merged ONT and merged HiFi assemblies against the 'Heinz 1706' SL4.0 assembly. Min identity (abs) was set to 0.65. **a,** alignment of merged NECAT+Flye to SL4.0. **b,** alignment of merged Hifiasm+-Canu to SL4.0. Assemblies have different breakpoints.

### Supplementary Figure 3

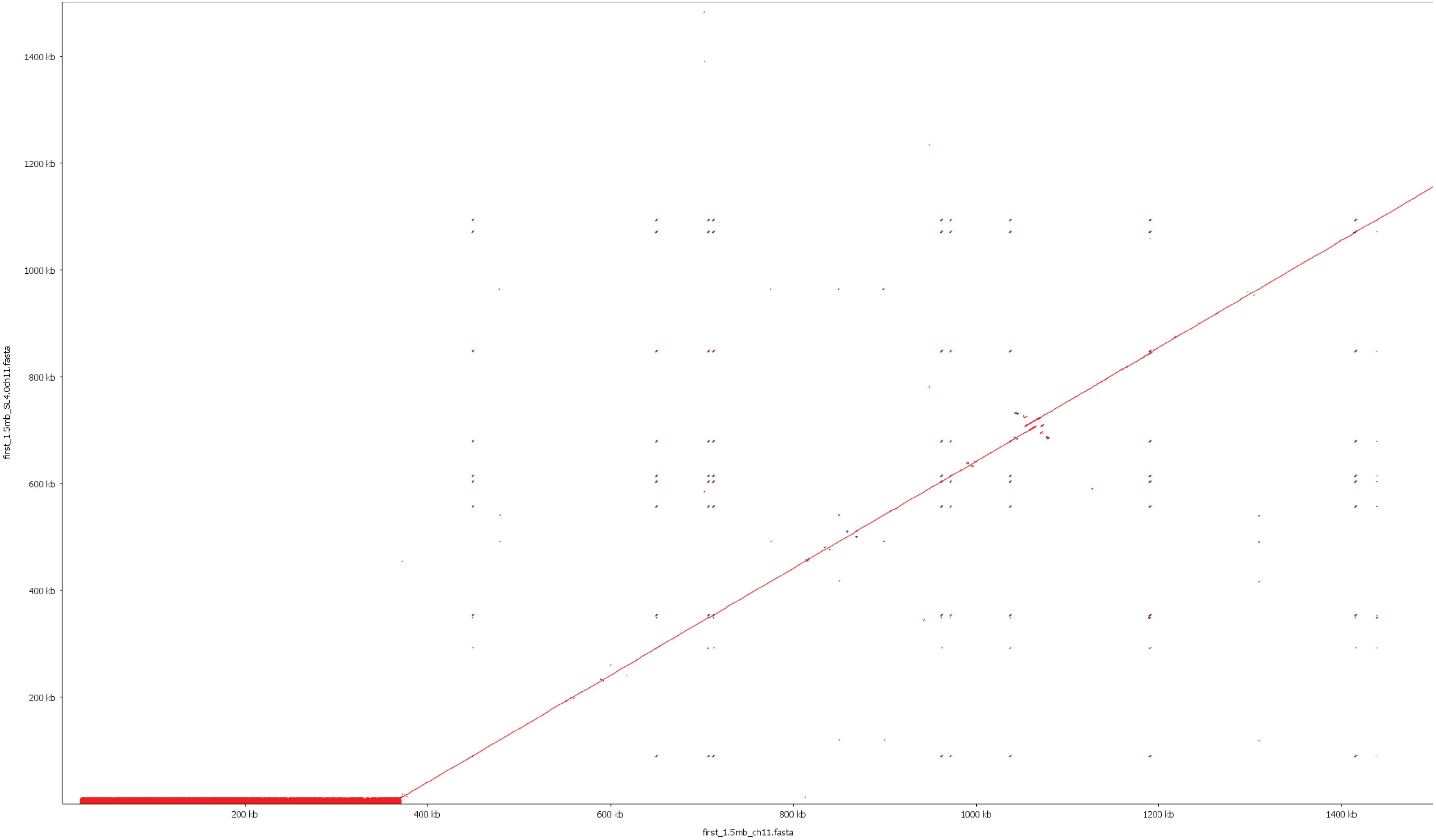


**Supplementary Fig. 3 | Alignment of the first 1.5 Mbp of MbTMV chromosome 11 with the first 1.5 Mbp of SL4.0 chromosome 11**
Alignment of first 1.5 Mbp of MbTMV ch11 (x-axis) with the first 1.5 Mbp of SL4.0 ch11 (y-axis). The SL4.0 sequence (~20 Kbp to ~1.1 Mbp) aligns near perfectly to MbTMV (~390 Kbp to 1.5 Mbp). The distal end of the MbTMV chromosome 11 (TGR1 dense region) is missing in the SL4.0 assembly.

### Supplementary Figure 4

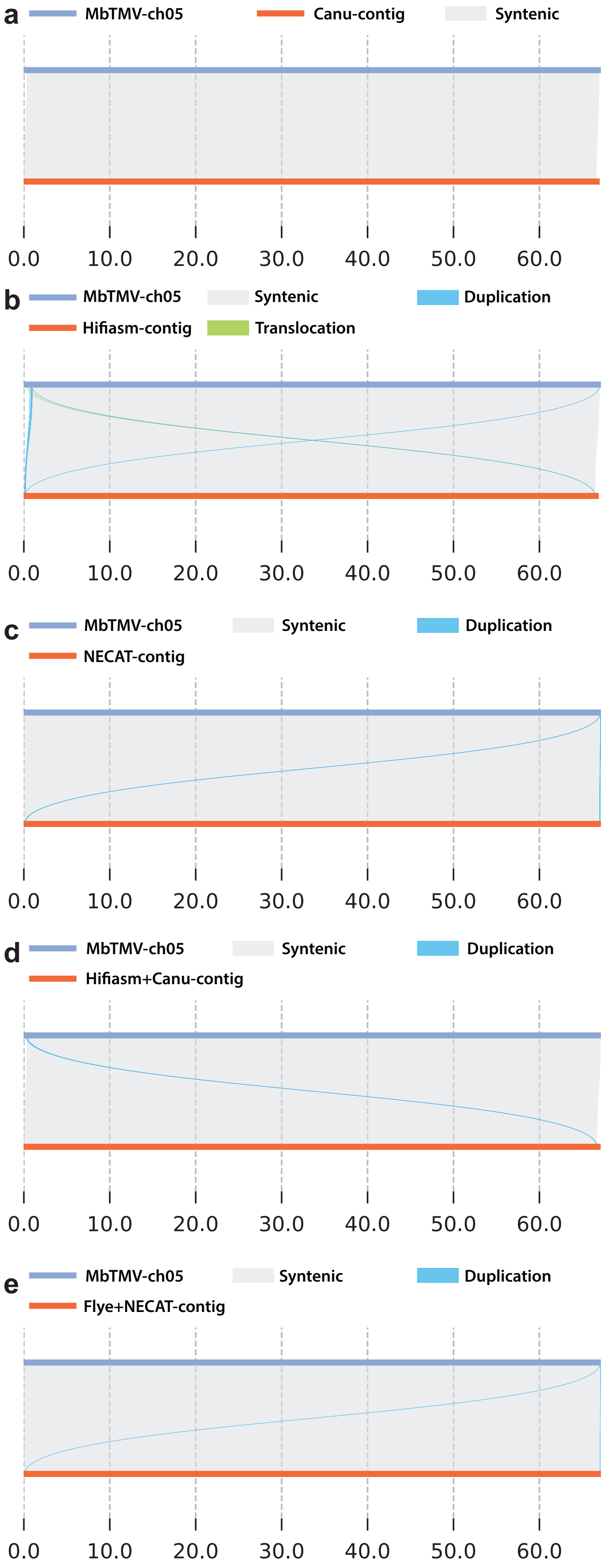


**Supplementary Fig. 4 | Synteny plots of different chromosome 5 assemblies**
SyRi plots (Goel et al., 2019) of single contigs, generated independently using various assembly tools, mapped against MbTMV-ch05. X-axis represents position on MbTMV-ch05 in megabasepair. All contigs span the centromere located between roughly 28 and 30 Mbp (Fig. 2e and Fig. 2f) showing only high synteny. **a**, Canu assembly contig “tig00000001” vs MbTMV-ch05. **b**, Hifiasm assembly contig “ptg000041l” vs MbTMV-ch05. **c**, NECAT assembly contig “bctg00000002” vs MbTMV-ch05. **d**, Hifiasm+Canu merged assembly contig “tig00000001” vs MbTMV-ch05. **e**, Flye+NECAT merged assembly contig “bctg00000002” vs MbTMV-ch05.

### Supplementary Figure 5

a

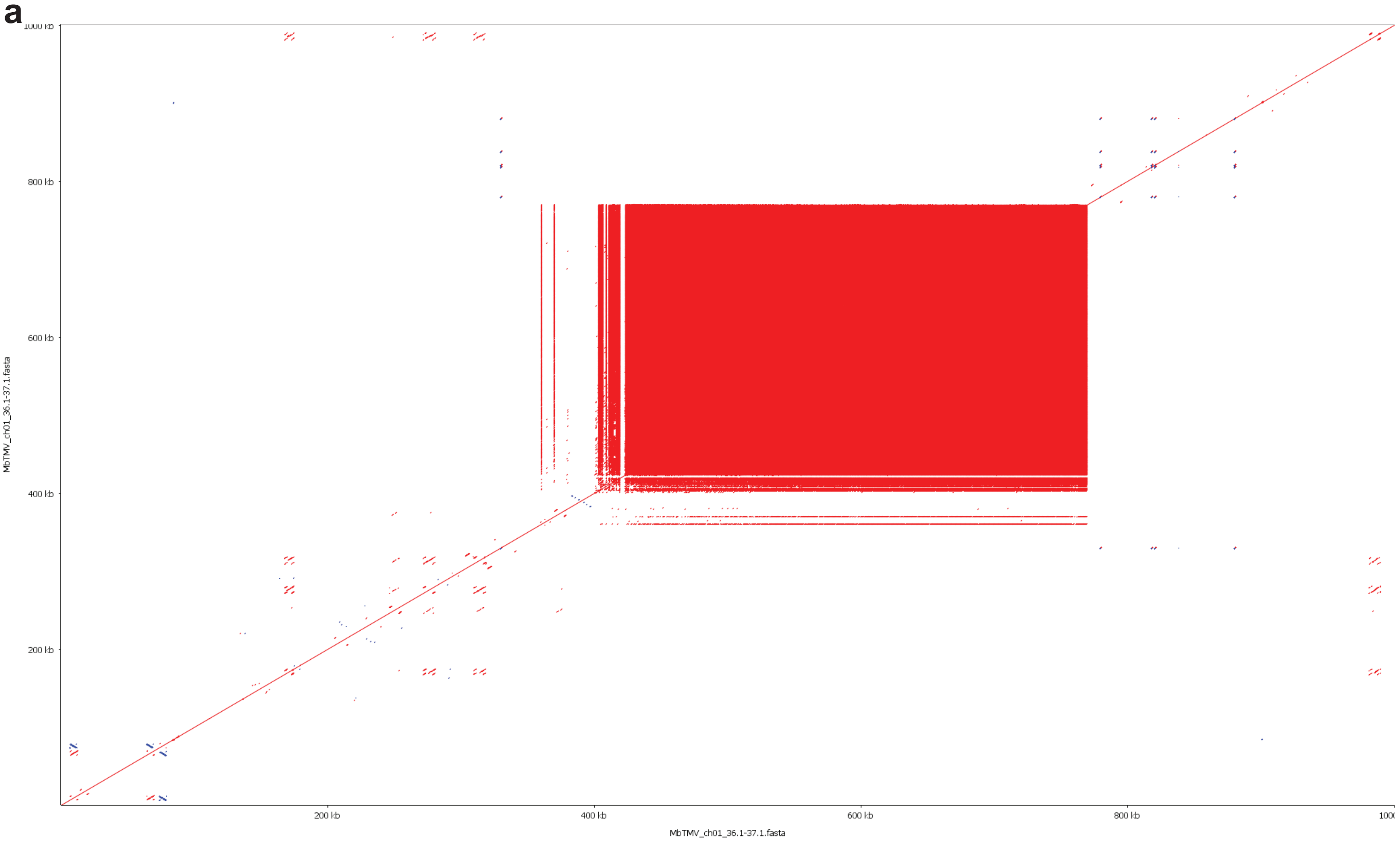


b

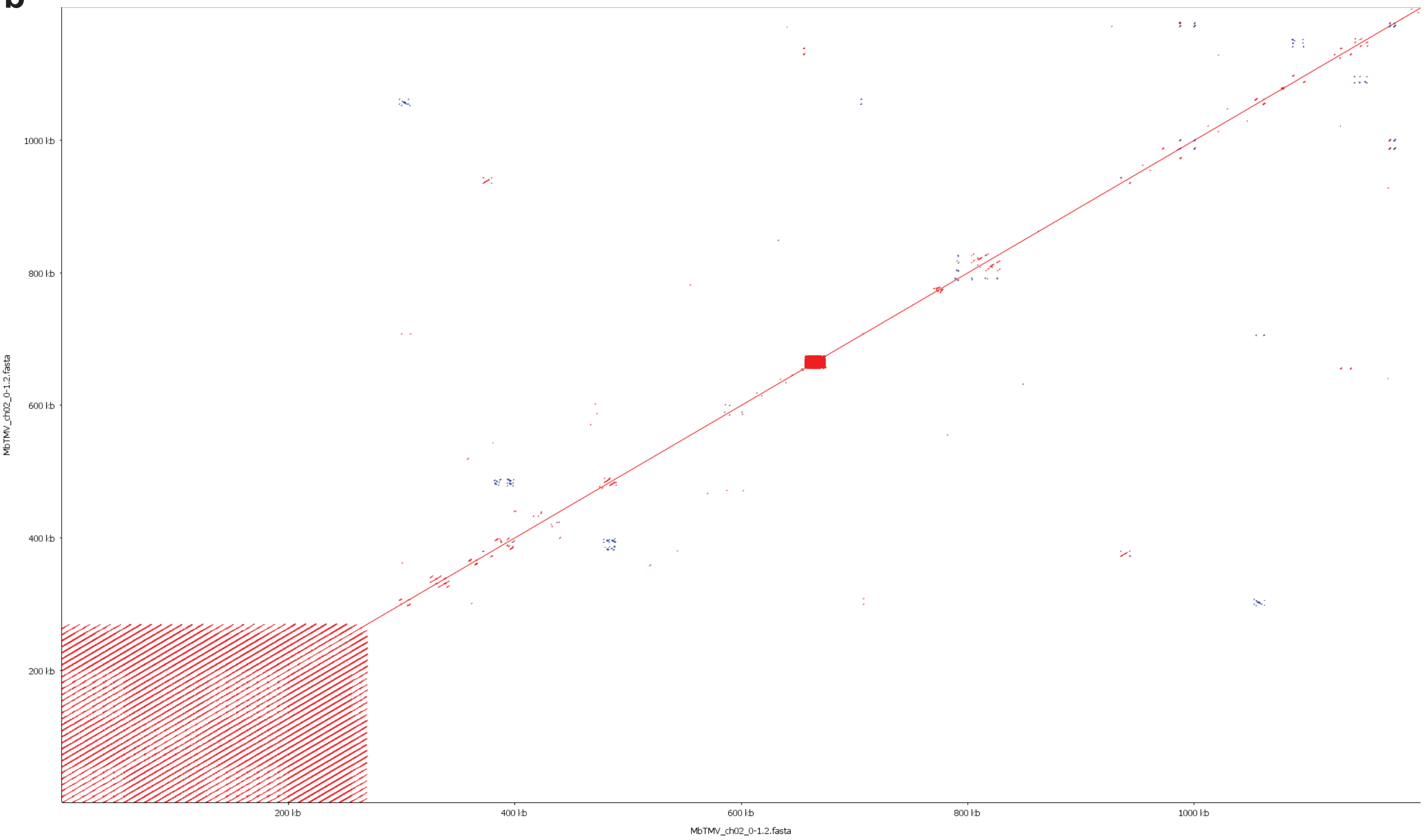


**Supplementary Fig. 5 | Self alignment plots of repetitive regions on MbTMV chromosomes 1 and 2**
**a**, Self alignment of a 1.2 Mbp region (36.1 Mbp to 37.1 Mbp) of MbTMV chromosome 1, showing a tandemly repeated region (5S rDNA) matching the coverage peak in Fig 2b. **b**, Self alignment a 1.2 Mbp region (1 bp to 1.2 Mbp) of MbTMV chromosome 2, showing a tandemly repeated region (45S rDNA) matching the coverage peak in Fig 2b.

### Supplementary Figure 6

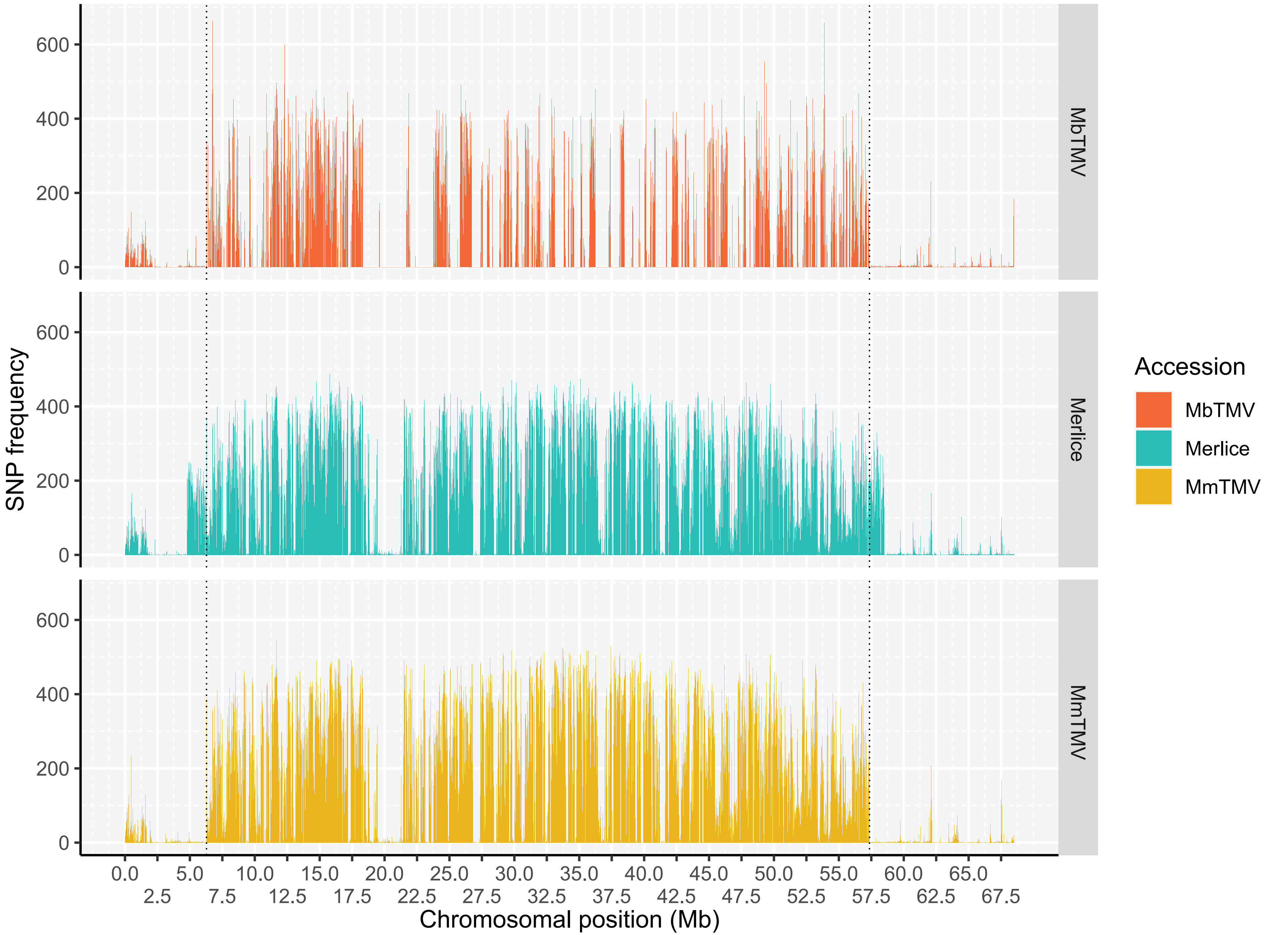


**Supplementary Fig. 6 | SNP frequency plot between MbTMV and SL4.0 in 10 kbp windows along chromosome 9**

MbTMV, MmTMV (Moneymaker-TMV) and Merlice variant calling against SL4.0. SNP frequency (calculated in a 10kb window), plotted over the length of SL4.0 chromosome 9. The *S. peruvianum* introgression was identified by increased SNP frequency. The two dotted vertical lines indicate identical introgression start and endpoints for MbTMV and MmTMV

### Supplementary Figure 7

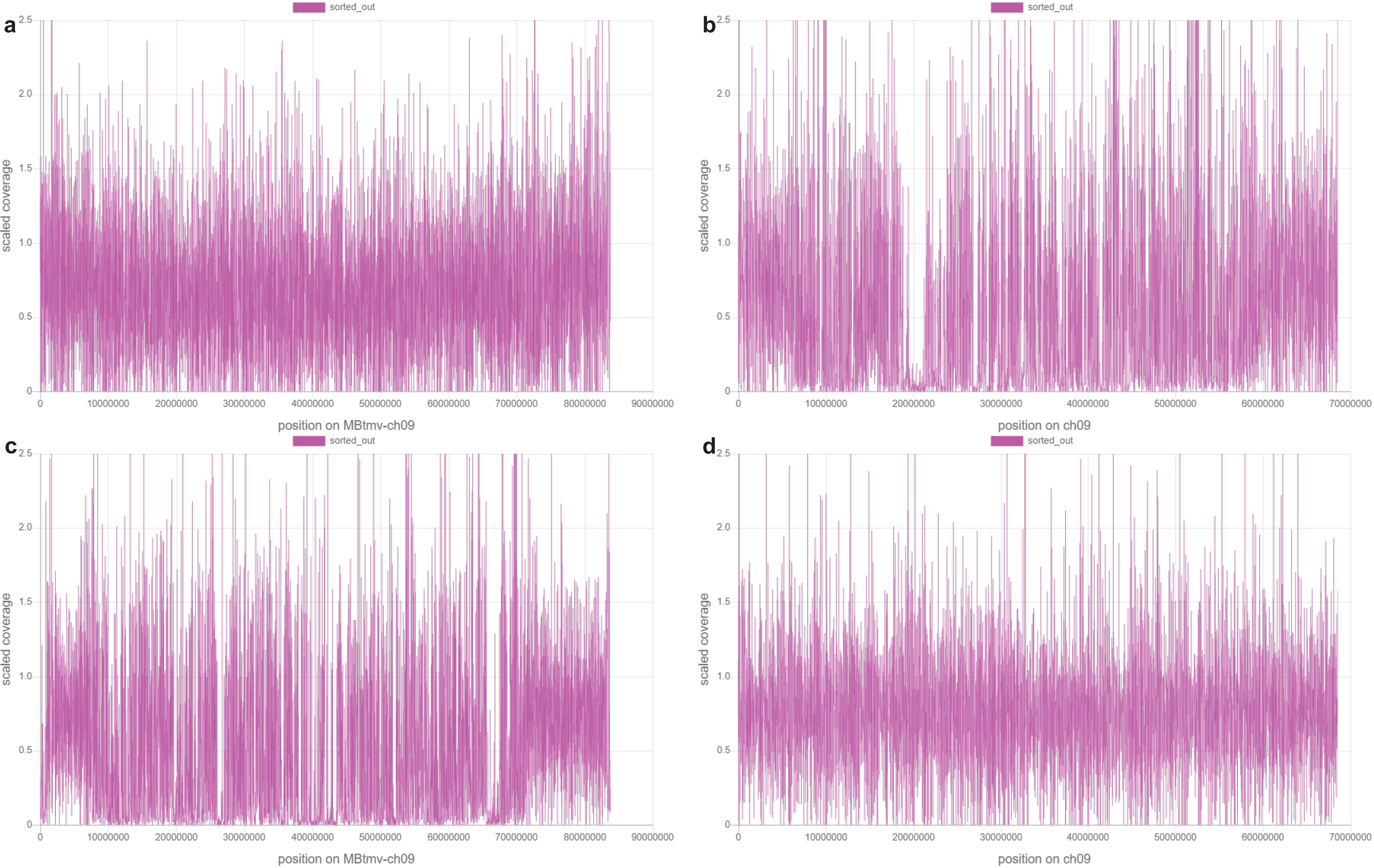


**Supplementary Fig. 7 | Coverage plot of LYC1969 and M82 ONT reads aligned to MbTMV chromosome 9 and SL4.0 chromosome 9**

Scaled coverage plot of LYC1969 and M82 ONT data (Alonge et al., 2020) aligned to chromosome 9 of MbTMV and SL4.0. Plots created with goleft IndexCov. **a**, LYC1969 ONT reads aligned to MbTMV. **b**, LYC1969 ONT reads aligned to SL4.0. **c**, M82 ONT reads aligned to MbTMV. **d**, M82 ONT reads aligned to SL4.0. A clear drop in coverage is shown for LYC1969 reads mapped to SL4.0 and M82 reads mapped to MbTMV.

### Supplementary Figure 8

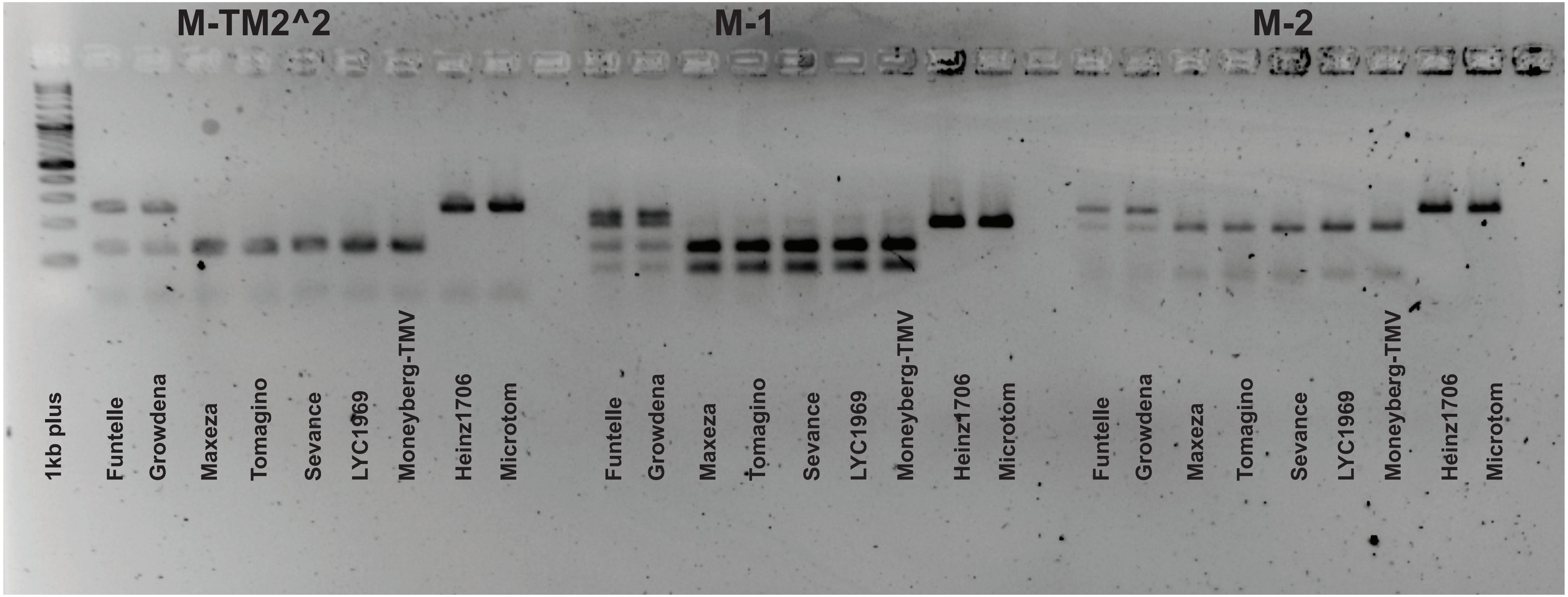


**Supplementary Fig. 8 | Agarose gel image of molecular markers**
Agarose gel electrophoresis of molecular markers M-1, M-TM2^2 and M-2 on 9 *S. lycopersicum* varieties including 5 commercial hybrid lines (Funtelle, Growdena, Maxeza, Tomagino and Sevance), 2 inbred lines resistant to TMV (LYC1969 and Moneyberg-TMV) and 2 inbred lines susceptible to TMV (Heinz1706 and Microtom). M-1 was digested with AflII (NEB), M-TM2^2 was digested with BslI (NEB) and M-2 was digested with XhoI (NEB). DNA ladder is 1kb plus ladder (NEB).

### Supplementary Figure 9

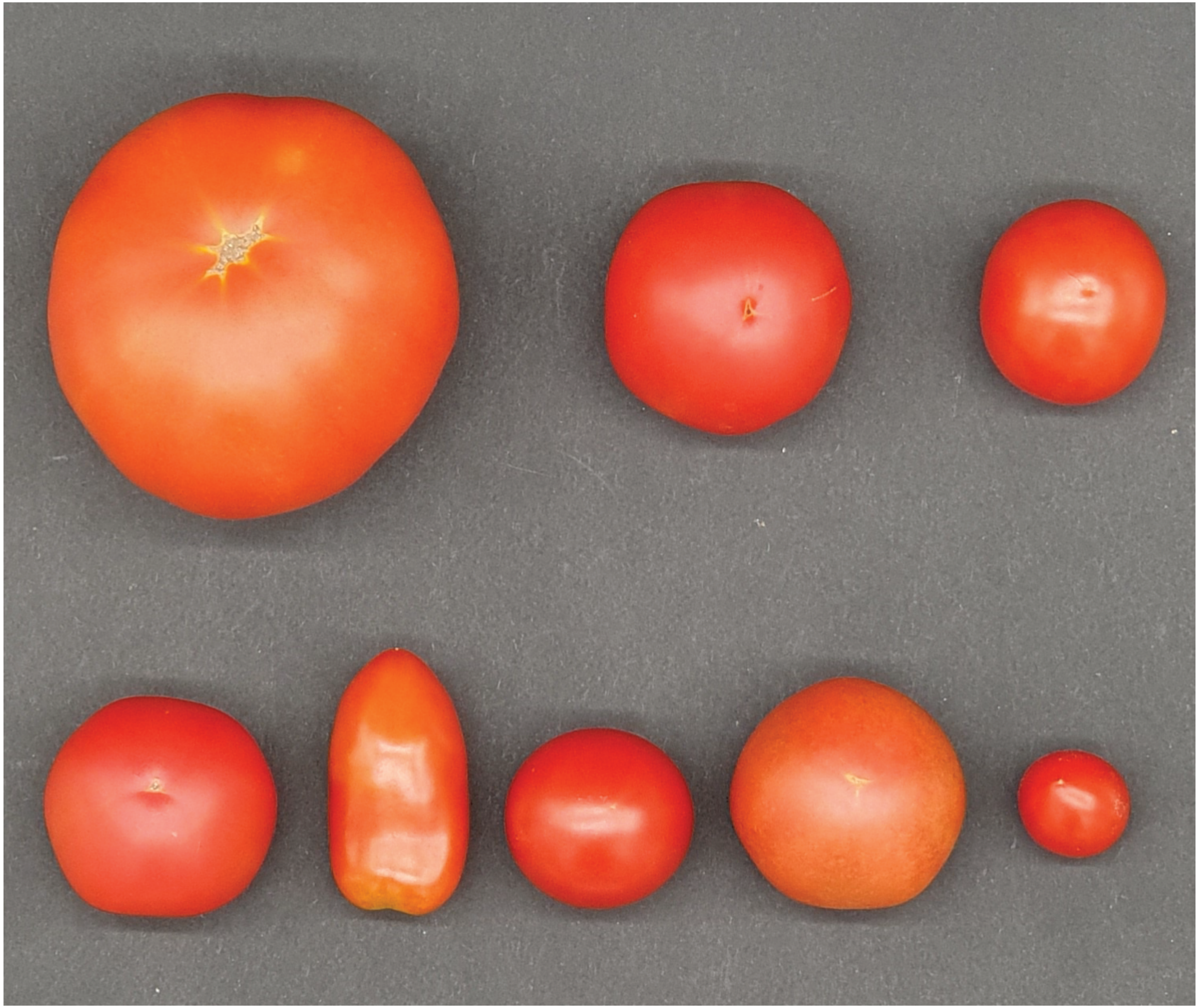

**Supplementary Fig. 9 | Fruit phenotypes of tomato varieties used in this study**
Top row, from left: Growdena, Maxeza, Tomagino; Bottom row, from left: Sevance, Funtelle, Moneyberg-TMV x Microtom F1 hybrid, Moneyberg-TMV, Microtom.
