## Supplementary Tables for "A gap-free tomato genome built from complementary PacBio and Nanopore long DNA sequences reveals extensive linkage drag during breeding"

### Supplementary Table 1) Statistics of different scaffolded assemblies

Supplementary table 1 shows the summary statistics for RaGOO scaffolded assemblies. BUSCO analysis of the completeness of gene content is using the *Solanales* benchmark set.

| Assembly | Hifiasm | Canu | NECAT | Flye | NECAT + Flye | Hifiasm + Canu | MbTMV |
|---|---|---|---|---|---|---|---|
| Scaffolding | RaGOO | RaGOO | RaGOO | RaGOO | RaGOO | RaGOO | RaGOO |
| Number of scaffolds | 14 | 14 | 14 | 14 | 14 | 14 | 13 |
| Cumulative size (Mbp) | 869 | 928.9 | 829.6 | 809.8 | 829 | 871.9 | 833 |
| N50 (Mbp) | 68.4 | 69 | 68.1 | 66.2 | 67.9 | 68.6 | 68.5 |
| N90 (Mbp) | 52.1 | 55.5 | 55.2 | 54.3 | 54.7 | 52.3 | 54.7 |
| L50 | 6 | 6 | 6 | 6 | 6 | 6 | 6 |
| L90 | 12 | 12 | 11 | 11 | 11 | 12 | 11 |
| Maximum scaffold size (Mbp) | 96.5 | 102.1 | 97.3 | 96.7 | 97.1 | 96.6 | 96.5 |
| Ns per 100 kbp | 8.47 | 25.64 | 1.34 | 4.68 | 1.11 | 8.22 | 0.64 |
| BUSCO (genes searched) | 5950 | 5950 | 5950 | 5950 | 5950 | 5950 | 5950 |
| BUSCO (complete) | 5853 | 5853 | 5779 | 5818 | 5823 | 5853 | 5853 |
| BUSCO (complete single) | 5743 | 5743 | 5674 | 5712 | 5715 | 5743 | 5743 |
| BUSCO (complete duplicated) | 110 | 110 | 105 | 106 | 108 | 110 | 110 |
| BUSCO (fragmented) | 12 | 12 | 38 | 21 | 17 | 12 | 12 |
| BUSCO (missing) | 85 | 85 | 133 | 111 | 110 | 85 | 85 |

**Supplementary Table 2) BUSCO results of different scaffolded assemblies using eudicot dataset**

Supplementary table 2 shows the BUSCO results, describing the completeness of gene content, of merged assemblies and established genomes mentioned in table 3, using the *Eudicots* benchmark set.

| Assembly name | MbTMV | NECAT + Flye | Hifiasm + Canu | SL4.0 | LA2093 | LA0716 |
|---|---|---|---|---|---|---|
| BUSCO (genes searched) | 2326 | 2326 | 2326 | 2326 | 2326 | 2326 |
| BUSCO (complete) | 2298 | 2291 | 2297 | 2290 | 2295 | 2288 |
| BUSCO (complete single) | 2276 | 2268 | 2276 | 2270 | 2278 | 2260 |
| BUSCO (complete duplicated) | 22 | 23 | 21 | 20 | 17 | 28 |
| BUSCO (fragmented) | 10 | 12 | 10 | 15 | 10 | 17 |
| BUSCO (missing) | 18 | 23 | 19 | 21 | 21 | 21 |

**Supplementary Table 3) Downsampling of PacBio HiFi reads and assembly using Hifiasm**

Supplementary table 3 shows the summary statistics of Hifiasm assemblies after downsampling of PacBio HiFi reads.

| Assembly | 100% | 90% | 80% | 70% | 60% | 50% | 40% | 30% | 20% | 10% |
|---|---|---|---|---|---|---|---|---|---|---|
| Number of contigs | 750 | 713 | 729 | 701 | 704 | 681 | 630 | 648 | 1260 | 6522 |
| Cumulative size (Mbp) | 868.9 | 865.5 | 862.3 | 860.3 | 858.2 | 854.3 | 848.1 | 842.1 | 825.7 | 672.3 |
| N50 (Mbp) | 31.3 | 31.3 | 31.3 | 31.3 | 20.8 | 20.1 | 18.6 | 9.0 | 1.5 | 0.1 |
| N90 (Mbp) | 8.7 | 7.3 | 6.7 | 6.6 | 6.6 | 7.2 | 5.3 | 2.6 | 0.4 | 0.1 |
| L50 | 10 | 10 | 10 | 10 | 11 | 14 | 14 | 31 | 162 | 1666 |
| L90 | 32 | 34 | 35 | 35 | 38 | 41 | 45 | 91 | 563 | 4865 |
| Longest contig (Mbp) | 66.6 | 60.9 | 66.0 | 66.0 | 66.3 | 52.2 | 52.5 | 40.6 | 8.0 | 0.7 |

**Supplementary Table 4) Downsampling of ONT reads and assembly using NECAT**

Supplementary table 4 shows the summary statistics of NECAT assemblies after downsampling of ONT reads. For ONT genome assemblies only ONT reads with an average Q-value of 9 or higher (in total 165.7 Gbp) were used. Large assembly artifacts were identified by comparison with non-downsampled Flye, Hifiasm and Canu assemblies.

| Assembly | 100% | 90% | 80% | 70% | 60% | 50% | 40% | 30% | 20% | 10% |
|---|---|---|---|---|---|---|---|---|---|---|
| Number of contigs | 125 | 110 | 140 | 131 | 126 | 122 | 138 | 143 | 229 | 418 |
| Cumulative size (Mbp) | 829.5 | 825.3 | 831.0 | 828.9 | 830.5 | 827.2 | 827.4 | 829.0 | 827.6 | 823.5 |
| N50 (Mbp) | 50.2 | 37.6 | 37.7 | 47.4 | 51.1 | 37.4 | 37.7 | 37.5 | 19.2 | 7.2 |
| N90 (Mbp) | 10.6 | 13.3 | 8.7 | 10.8 | 10.2 | 10.0 | 10.2 | 10.1 | 7.0 | 2.4 |
| L50 | 7 | 7 | 8 | 7 | 7 | 8 | 8 | 8 | 11 | 37 |
| L90 | 13 | 21 | 21 | 20 | 21 | 22 | 24 | 23 | 39 | 109 |
| Longest contig (Mbp) | 74.2 | 74.3 | 69.2 | 85.7 | 74.9 | 68.9 | 69.0 | 69.0 | 65.8 | 19.8 |
| Large assembly artifacts | no | yes | no | yes | no | no | no | no | yes | yes |

**Supplementary Table 5) Variants called between MbTMV and SL4.0 on chromosome 9 from a whole genome alignment**

Supplementary table 5 (vcf file) shows SyRI called variants between MbTMV (Alt / Query) chromosome 9 and SL4.0 (Ref) chromosome 9, extracted from the whole genome alignment.

**Supplementary Table 6) Variants called between MbTMV and SL4.0 on chromosome 9 from a single chromosome alignment**

Supplementary table 6 (vcf file) shows SyRI called variants between MbTMV (Alt / Query) chromosome 9 and SL4.0 (Ref) chromosome 9, from a direct alignment of MbTMV chromosome 9 and SL4.0 chromosome 9.

**Supplementary Table 7) Variants called between MbTMV and SL4.0 in a ~10 Mbp extracted region of chromosome 9**

Supplementary table 7 (vcf file) shows SyRI called variants between MbTMV (Alt / Query) chromosome 9 and SL4.0 (Ref) chromosome 9, from an extracted region.

**Supplementary Table 8) Variants called between MbTMV and SL4.0 in a ~1 Mbp extracted region of chromosome 9**

Supplementary table 8 (vcf file) shows SyRI called variants between MbTMV (Alt / Query) chromosome 9 and SL4.0 (Ref) chromosome 9, from an extracted region.
